# In silico characterization of lysis and host-recognition modules in *Staphylococcus aureus* bacteriophage genomes

**DOI:** 10.64898/2026.06.13.732091

**Authors:** Ivan Armadi Hasugian, Na’ilah Insani Alifiyah

**Affiliations:** Department of Biology, Faculty of Science and Technology, Universitas Terbuka, Tangerang Selatan, Indonesia

**Keywords:** Bacteriophage, endolysin, host recognition, comparative genomics, *Staphylococcus aureus*

## Abstract

**Background/aim:** Antimicrobial resistance in methicillin-resistant *Staphylococcus aureus* (MRSA) requires precision non-antibiotic therapeutics, yet phage lytic efficacy is poorly predicted by phenotypic assays, as shown by paradoxical biofilm responses. This study characterized the genomic architecture of lytic *S. aureus* bacteriophages, focusing on the conservation of the lysis module and the variability of host-recognition modules, to provide a rational basis for phage candidate selection.

**Materials and methods:** Twenty-two complete *S. aureus* phage genomes were retrieved from NCBI GenBank. Genomic features were extracted with custom Biopython scripts. Lysis (endolysin, holin) and host-recognition (tail fiber/receptor-binding protein) modules were annotated and validated by InterPro domain analysis, with disrupted endolysins resolved by tBLASTn. Phylogeny was reconstructed from large terminase subunit (TerL) sequences using maximum likelihood.

**Results:** Genome size spanned three classes, from 17.5 to 148.6 kb. The LysK-type endolysin (CHAP–Amidase–SH3b) was highly conserved, whereas tail fiber/RBP genes were detected in only 14 of 22 phages. Domain analysis reclassified two proteins annotated as endolysins as virion-associated peptidoglycan hydrolases, and identified two independent mechanisms—HNH endonuclease insertion and intron splitting—that interrupt lysis-module genes and confound automated annotation. Maximum likelihood analysis recovered a strongly supported, highly conserved core clade with EW and SA13 as divergent lineages.

**Conclusion:** Lysis modules are conserved whereas host-recognition modules are variable, indicating that host recognition rather than the lytic enzyme is the principal determinant of host range and the more rational target for phage selection and engineering.

## 1. Introduction

Antimicrobial resistance (AMR) driven by methicillin-resistant *Staphylococcus aureus* (MRSA) has escalated into a major global health threat, intensifying the need for precision non-antibiotic therapeutics (World Health Organization, 2022; Habboush and Guzman, 2023). The structural resilience of MRSA, mediated by the *mecA* gene product PBP2a and by OatA-dependent peptidoglycan O-acetylation, undermines the efficacy of both conventional and last-line antibiotics (Jones et al., 2020; Sari et al., 2024). Lytic bacteriophages offer a rational alternative, delivering precise bactericidal activity through their lysis cassette—the holin and endolysin proteins that degrade peptidoglycan from within the host cell—without disrupting the commensal microbiota (Brüser and Mehner-Breitfeld, 2022; Liu et al., 2024).

Despite the conceptual elegance of phage lysis, *in vitro* phenotypic evaluation frequently yields unpredictable outcomes owing to complex biological barriers (Duarte et al., 2021; Anyaegbunam et al., 2022). A paradoxical observation illustrates this limitation: phage vB_SauP-436A markedly reduced MRSA biofilm biomass, whereas phage vB_SauM-515A1 increased biofilm biomass, even though both phages exhibited high planktonic lytic efficiency *in vitro* (Abdraimova et al., 2024; Hasugian and Zikriyani, 2025). This divergence demonstrates that phenotypic lysis efficacy alone is an insufficient predictor of therapeutic success, particularly in chronic biofilm-associated infection (Molendijk et al., 2024).

Complete genome sequences of *S. aureus* bacteriophages are now abundantly available in public repositories such as NCBI GenBank, and most tailed phages have recently been reclassified within the class Caudoviricetes, including the families Herelleviridae and Rountreeviridae, superseding the earlier morphology-based grouping (Barylski et al., 2020). However, the textual product names in these deposited records are frequently inconsistent with the actual protein domain architecture, so a comparative analysis that re-examines the lysis and host-recognition modules against curated domain databases offers an efficient and reproducible means of clarifying their true functional content without requiring the isolation of new phages (Hatfull and Hendrix, 2011; Moller et al., 2022).

Although numerous *S. aureus* phage genomes are publicly available, a systematic comparison of lysis and host-recognition modules across therapeutically relevant lytic phages remains limited. Therefore, this study aimed to characterize the lysis and host-recognition modules of lytic *S. aureus* bacteriophages and to map candidate genomic features that may underlie inter-phage differences in therapeutic response. Given a dataset limited to publicly deposited sequences, the study was designed to be descriptive and comparative, providing a rational genomic foundation for the selection of natural phage candidates and to inform subsequent experimental investigation rather than to dictate clinical formulation

## 2. Materials and methods

### 2.1. Genome retrieval and dataset curation

Genome sequences were retrieved from the NCBI GenBank database^1^ between 18 and 30 May 2026 using the search query “Staphylococcus phage complete genome”, supplemented by targeted retrieval of the specific strains examined in a previous study (Hasugian and Zikriyani, 2025), and downloaded as GenBank flat files. The initial set was curated against predefined criteria. The inclusion criteria were (i) records annotated as complete genome sequences, (ii) a lytic lifestyle inferred from genome annotation and the absence of lysogeny-associated genes (integrase, excisionase, CI repressor), and (iii) *S. aureus* as the annotated host. The exclusion criteria were (i) duplicate or near-identical genome sequences, (ii) the presence of lysogeny-associated genes, and (iii) jumbo genomes exceeding 200 kb. During curation, Staphylococcus phage vB_SauP_EBHT (NC_055906) was excluded as essentially identical to Staphylococcus phage Portland (MT926124), and Staphylococcus phage MarsHill (MW248466) was excluded for exceeding 200 kb. Staphylococcus phage Sb1M_6168 (MN336262) and Sb1M_9832 (MN336263) were retained as representatives of natural intra-clade variation within the Sb-1 lineage. One record, Staphylococcus phage vB_SauM-515A1 (MN047438), carried a Draft/Partial status flag and was retained as a single, explicitly noted exception to the complete-genome criterion because all modules analysed here are fully annotated and the phage is of direct comparative relevance to the previous study. Host identity of all 22 retained records was manually confirmed against the GenBank source entries prior to analysis. The final dataset comprised 22 genomes.

### 2.2. Genomic feature extraction

Genome metadata were extracted using custom scripts written in Python 3.12.10 with Biopython 1.87 (Cock et al., 2009) and pandas 3.0.3. Using the Entrez and SeqIO modules, the scripts parsed each record to obtain genome size, GC content, the number of coding sequences (CDS), and the number of tRNA genes. Taxonomic ranks (class, family, subfamily) were resolved from the taxonomy field of each record by ICTV rank suffix (- viricetes, -viridae, -virinae); subfamilies placed by NCBI in no family (the Azeredovirinae phages EW and SA13) were recorded as family Unassigned. CDS counts are reported as deposited in GenBank; de novo re-annotation by independent gene callers may yield slightly different totals (for example, 236 versus 238 predicted features for MN047438). All extracted values were verified manually against the source records and are summarised in Table 1.

**Table 1.** General characteristics of the 22 lytic *Staphylococcus aureus* bacteriophage genomes.

| Phage | Accession | Class | Family | Subfamily | Genome size (bp) | GC (%) | CDS count | tRNA count | NCBI status |
| --- | --- | --- | --- | --- | --- | --- | --- | --- | --- |
| Staphylococcus phage A5W | EU418428.2 | Caudoviricetes | Herelleviridae | Twortvirinae | 145,542 | 30.42 | 232 | 4 | Complete |
| Staphylococcus phage qdsa002 | KY779849.1 | Caudoviricetes | Herelleviridae | Twortvirinae | 142,499 | 30.26 | 231 | 0 | Complete |
| Staphylococcus phage vB_SauM-fRuSau02 | MF398190.1 | Caudoviricetes | Herelleviridae | Twortvirinae | 148,464 | 30.22 | 236 | 4 | Complete |
| Staphylococcus phage vB_SavM_JYL01 | MH159197.1 | Caudoviricetes | Herelleviridae | Twortvirinae | 141,384 | 30.17 | 219 | 0 | Complete |
| Staphylococcus phage Maine | MN045228.1 | Caudoviricetes | Herelleviridae | Twortvirinae | 141,712 | 30.41 | 219 | 4 | Complete |
| Staphylococcus phage vB_SauM-515A1 | MN047438.1 | Caudoviricetes | Herelleviridae | Twortvirinae | 148,511 | 30.24 | 236 | 4 | Draft/Partial |
| Staphylococcus phage vB_SauP-436A1 | MN150710.1 | Caudoviricetes | Rountreeviridae | Rakietenvirinae | 18,028 | 29.34 | 21 | 0 | Complete |
| Staphylococcus phage Sb1_8383 | MN336261.1 | Caudoviricetes | Herelleviridae | Twortvirinae | 139,606 | 30.40 | 167 | 0 | Complete |
| Staphylococcus phage Sb1M_6168 | MN336262.1 | Caudoviricetes | Herelleviridae | Twortvirinae | 138,231 | 30.43 | 165 | 0 | Complete |
| Staphylococcus phage Sb1M_9832 | MN336263.1 | Caudoviricetes | Herelleviridae | Twortvirinae | 138,231 | 30.43 | 164 | 0 | Complete |
| Staphylococcus phage Portland | MT926124.1 | Caudoviricetes | Rountreeviridae | Rakietenvirinae | 17,471 | 29.57 | 20 | 0 | Complete |
| Staphylococcus phage Twort | NC_007021.1 | Caudoviricetes | Herelleviridae | Twortvirinae | 130,706 | 30.26 | 195 | 0 | Complete |
| Staphylococcus phage EW | NC_007056.1 | Caudoviricetes | Unassigned | Azeredovirinae | 45,286 | 35.99 | 77 | 0 | Complete |
| Staphylococcus phage JD007 | NC_019726.1 | Caudoviricetes | Herelleviridae | Twortvirinae | 141,836 | 30.37 | 217 | 0 | Complete |
| Staphylococcus phage SA13 | NC_021863.1 | Caudoviricetes | Unassigned | Azeredovirinae | 42,652 | 35.42 | 62 | 0 | Complete |
| *Staphylococcus aureus* bacteriophage Sb-1 | NC_023009.1 | Caudoviricetes | Herelleviridae | Twortvirinae | 127,188 | 30.48 | 168 | 4 | Complete |
| Staphylococcus phage Staph1N | NC_047722.1 | Caudoviricetes | Herelleviridae | Twortvirinae | 145,647 | 30.41 | 232 | 0 | Complete |
| Staphylococcus phage A3R | NC_047723.1 | Caudoviricetes | Herelleviridae | Twortvirinae | 141,018 | 30.50 | 212 | 0 | Complete |
| Staphylococcus phage 676Z | NC_047724.1 | Caudoviricetes | Herelleviridae | Twortvirinae | 148,564 | 30.41 | 233 | 0 | Complete |
| Staphylococcus phage Fi200W | NC_047725.1 | Caudoviricetes | Herelleviridae | Twortvirinae | 148,481 | 30.39 | 233 | 0 | Complete |
| Staphylococcus phage MSA6 | NC_047726.1 | Caudoviricetes | Herelleviridae | Twortvirinae | 148,243 | 30.24 | 232 | 0 | Complete |
| Staphylococcus phage P4W | NC_047727.1 | Caudoviricetes | Herelleviridae | Twortvirinae | 147,590 | 30.39 | 232 | 0 | Complete |

*“Unassigned” denotes a subfamily placed by NCBI/ICTV in no family (family incertae sedis), not a missing value.*

### 2.3. Functional annotation of lysis and host-recognition modules

Endolysin-candidate proteins were extracted from each genome and submitted to InterPro Version 108.0 (Blum et al., 2025) for domain architecture analysis against the Pfam, CDD, SUPERFAMILY, and Gene3D databases; this identified the CHAP, Amidase, SH3b, LysM, and NlpC/P60 domains reported in Table 2. Holin and tail fiber/receptor-binding protein (RBP) genes were detected from CDS product annotations using curated keyword sets, with each positive call retained together with the matched annotation for verification; the structural tail-tube protein, present in essentially all tailed phages, was deliberately excluded from the RBP keyword set. Where the automated candidate did not correspond to a recognized peptidoglycan hydrolase (Maine) or no intact endolysin-sized open reading frame was present (Sb1M_6168 and Sb1M_9832), the endolysin was treated as not confidently assigned by automated extraction; for the Sb-1 lineage the catalytic fragment was recovered by tBLASTn using the Sb1_8383 endolysin as query. In two further genomes, vB_SauM-515A1 (MN047438) and vB_SauM-fRuSau02 (MF398190), the endolysin (*lysK*) is annotated as one moiety because the gene is interrupted by a self-splicing intron; the intact enzyme is reconstituted at the RNA level, as demonstrated transcriptomically for vB_SauM- 515A1 (Kornienko et al., 2020), and the corresponding Table 2 entries are annotated accordingly.

**Table 2.** Domain architecture of lysis modules and presence of host-recognition modules among the 22 bacteriophages, validated by InterPro.

| Phage | Accession | Endolysin gene product | Endolysin length (aa) | Catalytic domains | Wall-binding domain | Holin present | Tail fiber/RBP present |
| --- | --- | --- | --- | --- | --- | --- | --- |
| Staphylococcus phage A5W | EU418428.2 | LysK | 495 | CHAP, Amidase | SH3b | Yes | Yes |
| Staphylococcus phage qdsa002 | KY779849.1 | Lysin | 495 | CHAP, Amidase | SH3b | Yes | No |
| Staphylococcus phage vB_SauM-fRuSau02 | MF398190.1 | LysK (intron-split) | 209 (moiety; full LysK ~495) | Amidase (moiety; CHAP intron-separated) | SH3b | Yes | Yes |
| Staphylococcus phage vB_SavM_JYL01 | MH159197.1 | Lysin | 495 | CHAP, Amidase | SH3b | Yes | Yes |
| Staphylococcus phage Maine | MN045228.1 | LysK-type endolysin (recovered by tBLASTn; missed by keyword extraction) | 495 (tBLASTn vs Sb1_8383, 99% identity) | CHAP, Amidase | SH3b | Yes | Yes |
| Staphylococcus phage vB_SauM-515A1 | MN047438.1 | LysK (intron-split) | 209 (moiety; full LysK ~495) | Amidase (moiety; CHAP intron-separated) | SH3b | Yes | Yes |
| Staphylococcus phage vB_SauP-436A1 | MN150710.1 | Endolysin (CHAP-type) | 250 | CHAP | SH3b | No | Yes |
| Staphylococcus phage Sb1_8383 | MN336261.1 | Putative endolysin | 494 | CHAP, Amidase | SH3b | Yes | No |
| Staphylococcus phage Sb1M_6168 | MN336262.1 | Endolysin (HNH-disrupted) | 141 (catalytic fragment, tBLASTn) | CHAP | LysM (separately annotated) | Yes (putative) | No |
| Staphylococcus phage Sb1M_9832 | MN336263.1 | Endolysin (HNH-disrupted) | 141 (catalytic fragment, tBLASTn) | CHAP | LysM (separately annotated) | Yes (putative) | No |
| Staphylococcus phage Portland | MT926124.1 | Lysin | 249 | CHAP | SH3b | Yes | Yes |
| Staphylococcus phage Twort | NC_007021.1 | Tail-associated lysin (VAPH) | 1,269 | Phage tail lysozyme (Phage_lysozyme2) | N/A (tail-anchored) | Yes | Yes |
| Staphylococcus phage EW | NC_007056.1 | Endolysin (divergent) | 482 | Non-LysK (divergent) | — | Yes | Yes |
| Staphylococcus phage JD007 | NC_019726.1 | LysK (recovered by tBLASTn; missed by keyword extraction) | 495 (tBLASTn vs Sb1_8383, 99% identity) | CHAP, Amidase | SH3b | Yes | Yes |
| Staphylococcus phage SA13 | NC_021863.1 | Endolysin | 481 | CHAP, Amidase | SH3b | Yes | Yes |
| *Staphylococcus aureus* bacteriophage Sb-1 | NC_023009.1 | Endolysin | 494 | CHAP, Amidase | SH3b | Yes | Yes |
| Staphylococcus phage Staph1N | NC_047722.1 | LysKn/LysKc | 495 | CHAP, Amidase | SH3b | Yes | Yes |
| Staphylococcus phage A3R | NC_047723.1 | LysK | 495 | CHAP, Amidase | SH3b | Yes | Yes |
| Staphylococcus phage 676Z | NC_047724.1 | LysK | 495 | CHAP, Amidase | SH3b | Yes | No |
| Staphylococcus phage Fi200W | NC_047725.1 | LysK | 495 | CHAP, Amidase | SH3b | Yes | No |
| Staphylococcus phage MSA6 | NC_047726.1 | LysK | 495 | CHAP, Amidase | SH3b | Yes | No |
| Staphylococcus phage P4W | NC_047727.1 | LysK | 495 | CHAP, Amidase | SH3b | Yes | No |

For vB\_SauM-fRuSau02 and vB\_SauM-515A1 the lysK gene is interrupted by a self-splicing intron carrying the ksaI
homing endonuclease; the 209-aa value is one moiety (Amidase + SH3b), and the intact LysK restores the CHAP–
Amidase–SH3b architecture (Kornienko et al., 2020). For Sb1M\_6168 and Sb1M\_9832 the endolysin open reading frame
is disrupted by an HNH endonuclease; the catalytic fragment was inferred by tBLASTn (Kornienko et al., 2023). For
Maine and JD007, automated extraction returned a non-lytic YbiA-like N-glycosidase and a virion-associated NlpC/P60
protein, respectively; the genuine LysK endolysins (495 aa, 99% identity to Sb1\_8383) were recovered by tBLASTn. For
Twort, tBLASTn detected only a partial, divergent LysK region (45% identity over 456 aa), and its principal hydrolase is
the virion-associated tail lysozyme. VAPH, virion-associated peptidoglycan hydrolase; RBP, receptor-binding protein.

### 2.4. Phylogenetic analysis

Phylogenetic relationships were inferred from the large terminase subunit (TerL), a conserved marker suitable for resolving relationships across phage families (Meier-Kolthoff and Göker, 2017; Dion et al., 2020). TerL sequences were extracted from each genome; an exact-match check was required to capture the abbreviated “Ter” product name used in the Kayvirus group. Staphylococcus phage Portland (MT926124) and vB_SauP-436A1 (MN150710) lacked an annotated TerL—consistent with their atypical small genome sizes (<20 kb)—and were excluded, leaving 20 sequences. Sequences were aligned with MAFFT using the L-INS-i strategy (Katoh et al., 2019). A maximum likelihood tree was constructed in MEGA 12.1.2 (Stecher et al., 2026) under the LG+G+I substitution model, with rate variation modelled by a discrete Gamma distribution and a proportion of invariant sites; node support was assessed with 1000 bootstrap replicates, gaps were treated by partial deletion with an 80% site-coverage cutoff, the tree was rooted on Staphylococcus phage EW (NC_007056) as the outgroup, and nodes with bootstrap support below 50% were collapsed.

## 3. Results and discussion

### 3.1. General genomic characteristics of lytic *Staphylococcus aureus* bacteriophages

Analysis of 22 complete bacteriophage genomes retrieved from NCBI GenBank revealed substantial genomic variation among candidate therapeutic phages targeting *S. aureus* (Table 1). Genome size ranged more than eight-fold, from 17,471 bp in Staphylococcus phage Portland (MT926124) to 148,564 bp in Staphylococcus phage 676Z (NC_047724). GC content varied within a narrower range, from 29.34% in vB_SauP-436A1 (MN150710) to 35.99% in phage EW (NC_007056), and the number of predicted CDS ranged from 20 in phage Portland to 236 in both vB_SauM-fRuSau02 (MF398190) and vB_SauM-515A1 (MN047438).

On the basis of genome size and CDS count, the phages fell into three size classes. The micro class (17–18 kb, 20–21 CDS) comprised Portland and vB_SauP-436A1, both assigned to the family Rountreeviridae. The medium class (42–45 kb, 62–77 CDS) comprised phages EW and SA13. The macro class (127–148 kb, 164–236 CDS) encompassed the majority of the dataset, including the Kayvirus group and the Sb-1 lineage. This stratification is consistent with established morphological descriptions, in which larger contractile-tailed (myovirus-type) phages correspond to the macro class and smaller phages to the micro class; these morphological descriptors are used here only informally, following the genome-based reclassification noted above. Macro-class phages possess sufficient coding capacity to accommodate additional engineering modules, an advantage absent in micro-class phages whose genomes approach the functional minimum (Pires et al., 2022).

GC content was tightly constrained across the dataset: 20 of the 22 phages exhibited GC content below 31%, close to but slightly below the approximately 32%–33% genomic GC of *S. aureus*. Two phages, EW (35.99%) and SA13 (35.42%), were clear outliers with GC content above the host average; this compositional divergence is consistent with their divergent placement in the phylogenetic analysis (Section 3.3, Figure 1).

**Figure 1.**
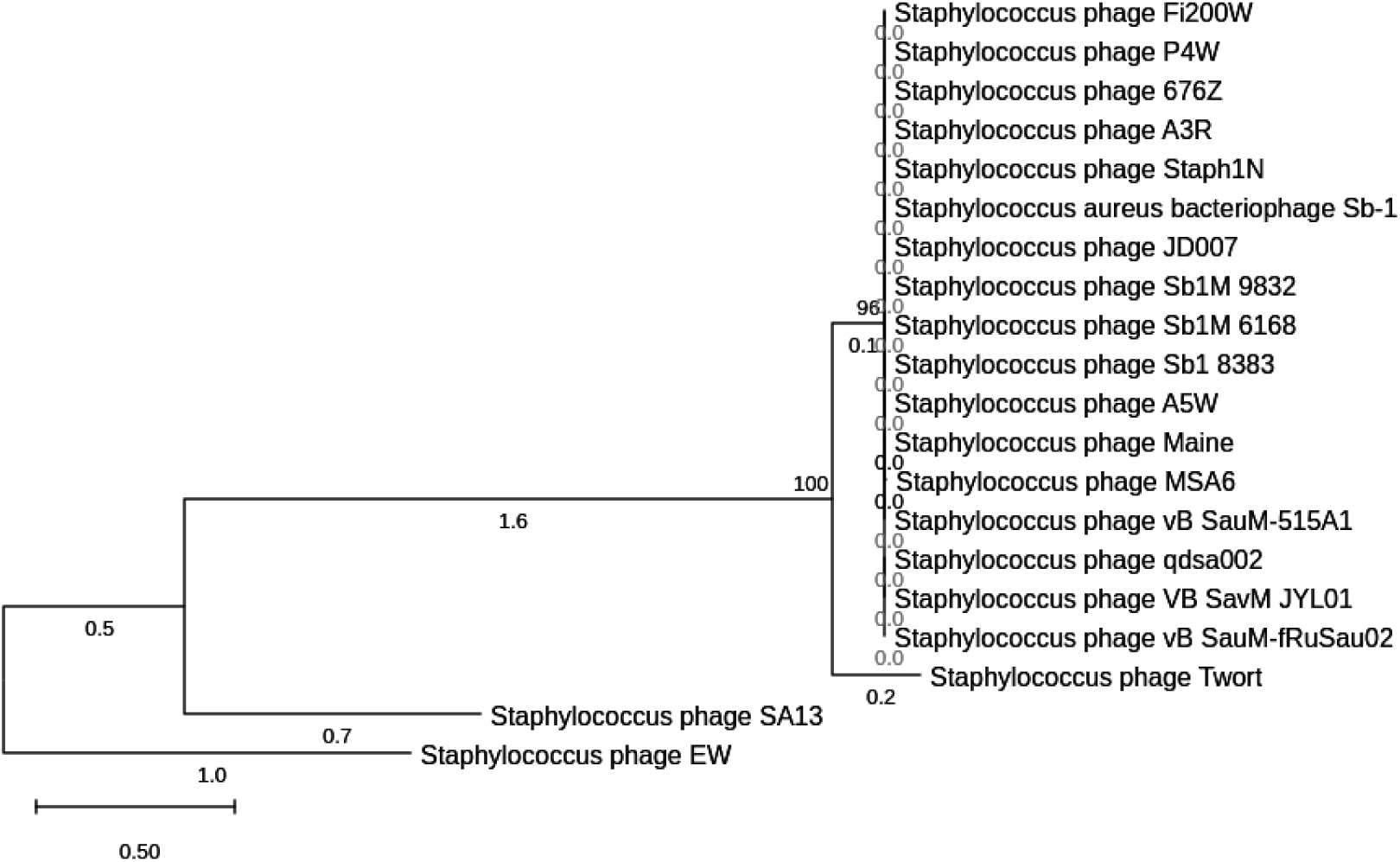
Maximum likelihood phylogenetic tree of 20 *Staphylococcus aureus* bacteriophages based on large terminase subunit (TerL) sequences, inferred in MEGA 12.1.2 under the LG+G+I model with 1000 bootstrap replicates and rooted on Staphylococcus phage EW (NC_007056). Bootstrap values are shown at the supported nodes; the scale bar indicates 0.50 substitutions per site.

Four tRNA genes were present in five phages—vB_SauM-fRuSau02, A5W, vB_SauM-515A1, Sb-1, and Maine—a genomic feature frequently overlooked in phenotypic analysis. Phage-encoded tRNAs have been proposed both as translational supplements that support phage protein synthesis when the host tRNA pool is suboptimal and, more recently, as anti-defense elements that counteract bacterial immune systems (Azam et al., 2024; Lomeli-Ortega and Balcázar, 2024; van den Berg and Brouns, 2025); both roles remain hypotheses requiring experimental confirmation in *S. aureus* phages.

Collectively, these features indicate that genomic characterization yields stratification information not obtainable from phenotypic data alone. The previous experimental evaluation of phage lytic activity by Hasugian and Zikriyani (2025) did not distinguish inter-phage genomic capacity as a variable influencing therapeutic outcome. The more than eleven-fold difference in CDS count between Portland (20 CDS) and vB_SauM-fRuSau02 (236 CDS) implies substantial differences in coding capacity for lysis strategy, accessory functions, and engineering potential, although the specific accessory-gene repertoires underlying these differences were not annotated in the present study.

### 3.2. Conservation of lysis modules and variability of host-recognition modules

Functional annotation of the 22 genomes revealed two biologically contrasting patterns: high conservation of the intracellular lysis components and substantial variability of the extracellular host-recognition components (Table 2). Among phages with a canonical free endolysin, protein length clustered tightly around 480–495 amino acids, and fourteen of these displayed the classic LysK-type architecture, defined by tandem N-terminal catalytic domains (CHAP and Amidase) and a C-terminal SH3b cell wall-binding domain (CBD); proteins falling outside this range corresponded to specific structural or genomic exceptions rather than to true size variation of the endolysin. Holin was detected in 21 of the 22 phages, whereas tail fiber or RBP genes were detected in only 14 of the 22, making host recognition the most heterogeneous feature in the dataset.

The LysK-type architecture acts cooperatively to cleave the pentaglycine cross-bridges characteristic of the *S. aureus* peptidoglycan (Gutiérrez et al., 2021). Its conservation extended even to the phylogenetically divergent phage SA13 (481 amino acids), which retained the full CHAP–Amidase–SH3b organisation despite its long branch and elevated GC content. The principal exception was phage EW, the most divergent lineage, whose 483-amino-acid endolysin diverges from the canonical LysK architecture, consistent with its phylogenetic position.

Two distinct departures from the canonical architecture were observed, both of which confound keyword-based annotation. In Sb1M_6168 and Sb1M_9832, the complete CDS inventory revealed no intact endolysin-sized open reading frame; instead, a 57-amino-acid LysM-domain protein and a 166-amino-acid putative HNH endonuclease were annotated separately within the lysis-module region. Standard BLASTp returned no significant similarity, whereas tBLASTn using the Sb1_8383 endolysin as query recovered a 141-amino-acid region homologous to LysK, whose CHAP domain was confirmed by InterPro. This pattern is consistent with HNH endonuclease insertion disrupting the continuity of the endolysin open reading frame in the Sb-1 lineage (Kornienko et al., 2023). A mechanistically different case was found in vB_SauM-515A1 and vB_SauM-fRuSau02, where extraction recovered only a 209-amino-acid moiety bearing the Amidase and SH3b domains but lacking the CHAP domain. Here the *lysK* gene is interrupted by a self-splicing group I intron carrying the *ksaI* homing endonuclease; the CHAP domain lies on the opposite side of the intron, and the intact LysK—restoring the canonical CHAP–Amidase–SH3b architecture—is reconstituted at the RNA level, as established transcriptomically for vB_SauM-515A1 (Kornienko et al., 2020). In both lineages, therefore, the apparently anomalous endolysin reflects interruption of the lysis-module gene by a mobile genetic element rather than a genuinely truncated enzyme, and both required manual curation rather than automated annotation alone.

In three further genomes, the automated keyword extractor returned a protein that was not the canonical free endolysin, and a targeted tBLASTn search using the Sb1_8383 endolysin as query was required to resolve each case. For Maine, the extractor recovered a 162-amino-acid YbiA-like N-glycosidase carrying no peptidoglycan-hydrolase domain, whereas tBLASTn recovered the genuine endolysin as a 495-amino-acid LysK (99% identity). For JD007, the extractor returned a 295-amino-acid protein combining an NlpC/P60 endopeptidase with a tail-associated Lid_Weld domain—a virion-associated peptidoglycan hydrolase—yet tBLASTn revealed that JD007 also encodes a separate, canonical 495-amino-acid LysK endolysin (99% identity) that had been overlooked. For Twort, the extractor returned a 1,269-amino-acid phage tail lysozyme (likewise a virion-associated hydrolase), and tBLASTn detected only a partial, divergent region (45% identity over 456 amino acids), indicating that Twort relies principally on its virion-associated tail lysozyme rather than on a canonical free endolysin. Together with the Sb-1 lineage, these cases show that keyword-based extraction can return non-lytic proteins, conflate virion-associated hydrolases with free endolysins, or miss the genuine enzyme altogether; reliable assignment therefore required domain validation by InterPro and homology search by tBLASTn rather than reliance on product names alone (Latka et al., 2017).

The contrast between a conserved lysis module and a variable host-recognition module indicates that efforts to overcome MRSA biofilm tolerance are more rationally directed at the host-recognition apparatus—tail fibers and RBPs—than at the internal lytic enzymes (Yehl et al., 2019; Pires et al., 2022). This interpretation bears directly on the paradoxical biofilm behaviour of vB_SauM-515A1 (RPCSa2) reported by Hasugian and Zikriyani (2025): because its endolysin is intron-split rather than truncated, and the intact LysK retains the full catalytic architecture, its atypical biofilm response is unlikely to stem from endolysin deficiency and is more plausibly related to its host-recognition characteristics.

### 3.3. Phylogenetic relationships based on the large terminase subunit

The maximum likelihood tree inferred from TerL sequences, rooted on phage EW, separated the 20 analysed phages into a strongly supported core clade and two divergent lineages (Figure 1). The core clade (bootstrap 100%) comprised 18 phages with large genomes (127–148 kb) consistent with a myovirus-type morphology; within it, Staphylococcus phage Twort (NC_007021) occupied the basal position, consistent with its recognised status as a prototype myovirus, while the remaining 17 phages formed a second supported group (bootstrap 96%) with near-zero internal branch lengths. Phages SA13 (NC_021863) and EW (NC_007056) formed the two divergent lineages, with long branches of 0.747 and 1.481 substitutions per site, respectively.

The near-zero internal branch lengths and consequent low internal support within the core clade reflect the high conservation of TerL among these phages rather than a methodological limitation, since a single conserved marker such as TerL resolves inter-family relationships well but discriminates poorly among closely related intra-family members (Meier-Kolthoff and Göker, 2017; Dion et al., 2020). The long branches and elevated GC content of EW and SA13 are mutually consistent and support independent evolutionary trajectories for these two phages. Importantly, the placement of vB_SauM- 515A1 within the conserved core clade indicates that its atypical biofilm behaviour cannot be attributed to deep phylogenetic divergence; combined with the conserved lysis architecture inferred for this phage (Table 2), this again points to determinants at the level of host recognition rather than lysis.

Several limitations should be acknowledged. The analysis is annotation-dependent: feature counts and module assignments derive from deposited GenBank records and computational domain inference, which approximate but do not replace experimental validation of protein function. The study is descriptive and comparative; no statistical hypothesis testing was applied, and the modest size of the dataset limits generalisation. The phylogeny rests on a single conserved marker and therefore does not capture the mosaic, recombination-rich nature of phage genomes. Accordingly, the genomic features identified here are best regarded as hypothesis-generating signals that require subsequent experimental confirmation, in line with the role of this study as preliminary groundwork for planned experimental investigation.

## 4. Conclusion

This comparative genomic survey of lytic *S. aureus* bacteriophages shows that the lysis module—endolysin and holin—is highly conserved, whereas host-recognition modules (tail fibers and RBPs) are markedly variable, being detectable in only 14 of the 22 genomes examined. Domain-level analysis distinguished true endolysins from virion-associated peptidoglycan hydrolases and revealed two independent mechanisms—HNH endonuclease insertion and intron splitting—by which lysis-module genes can be interrupted, both of which can mislead annotation based on product names alone. Because host recognition rather than the lytic enzyme emerges as the principal variable determinant of host range, it represents the more rational target for both candidate selection and engineering; experimental studies have likewise shown that host range can be broadened by modifying tail fiber proteins rather than the endolysin (Ando et al., 2015; Yehl et al., 2019). These findings provide a genomic foundation for the rational selection of phage candidates and a basis for the experimental investigation of planktonic and biofilm-associated responses in subsequent work.

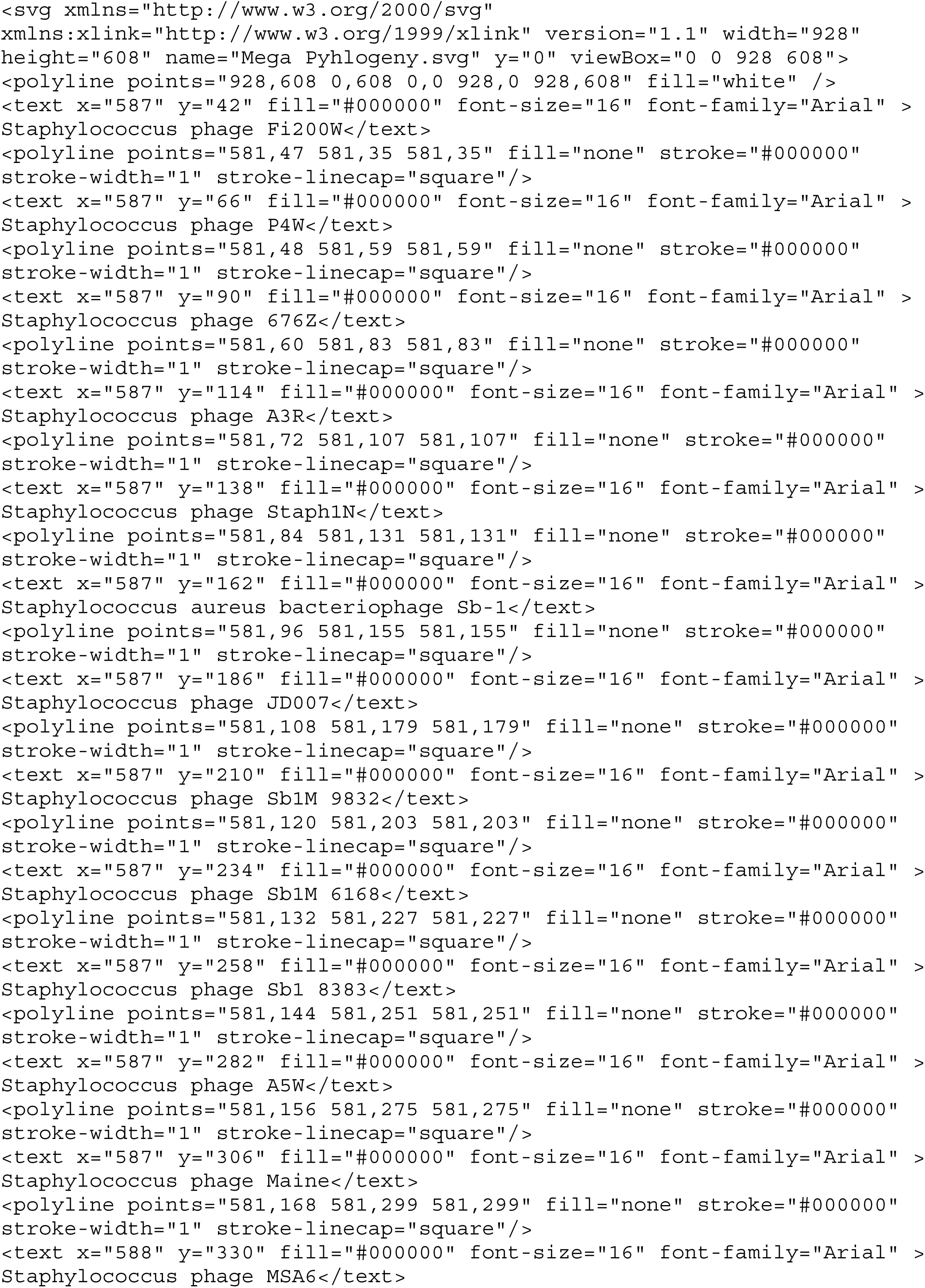
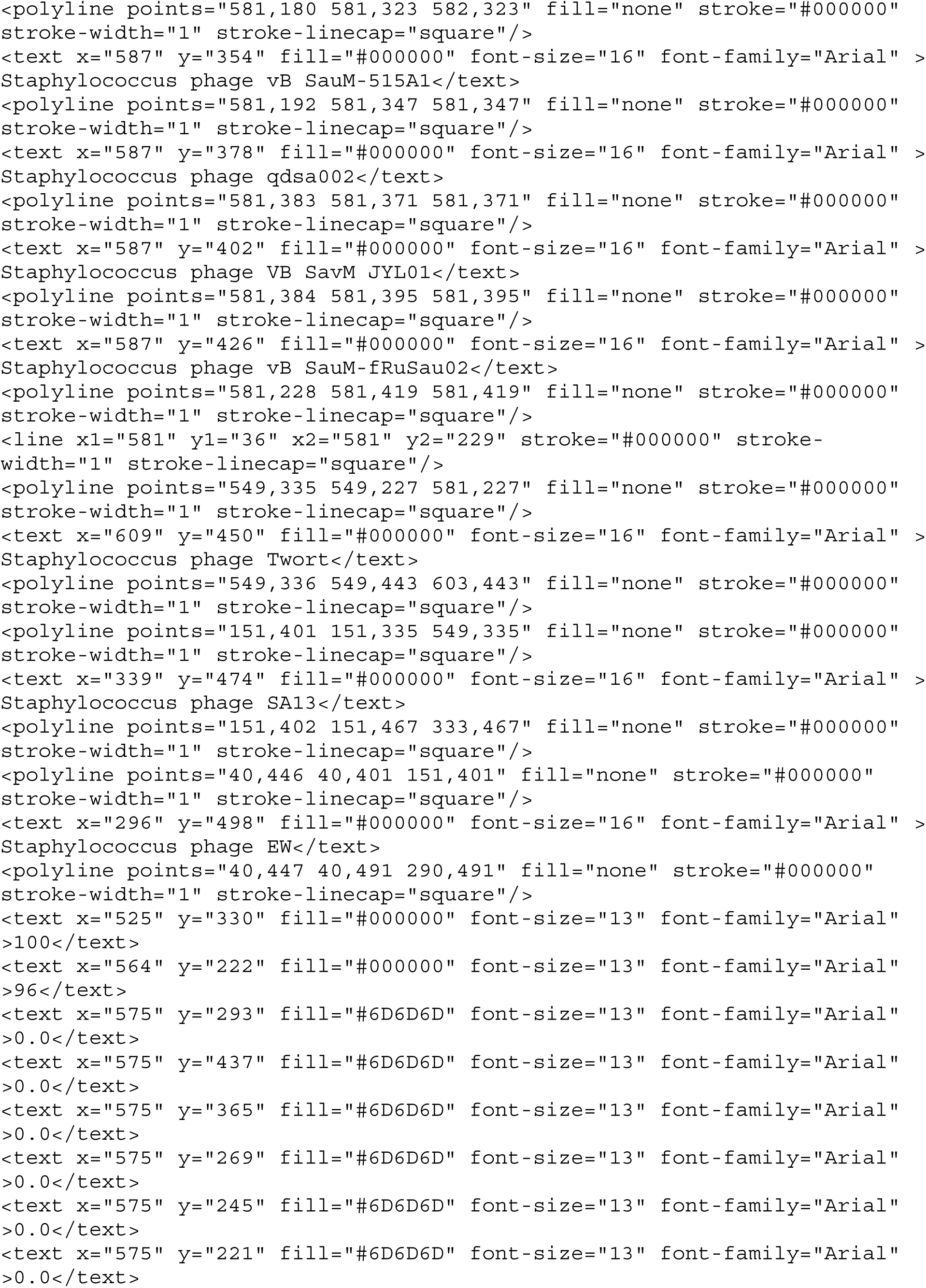
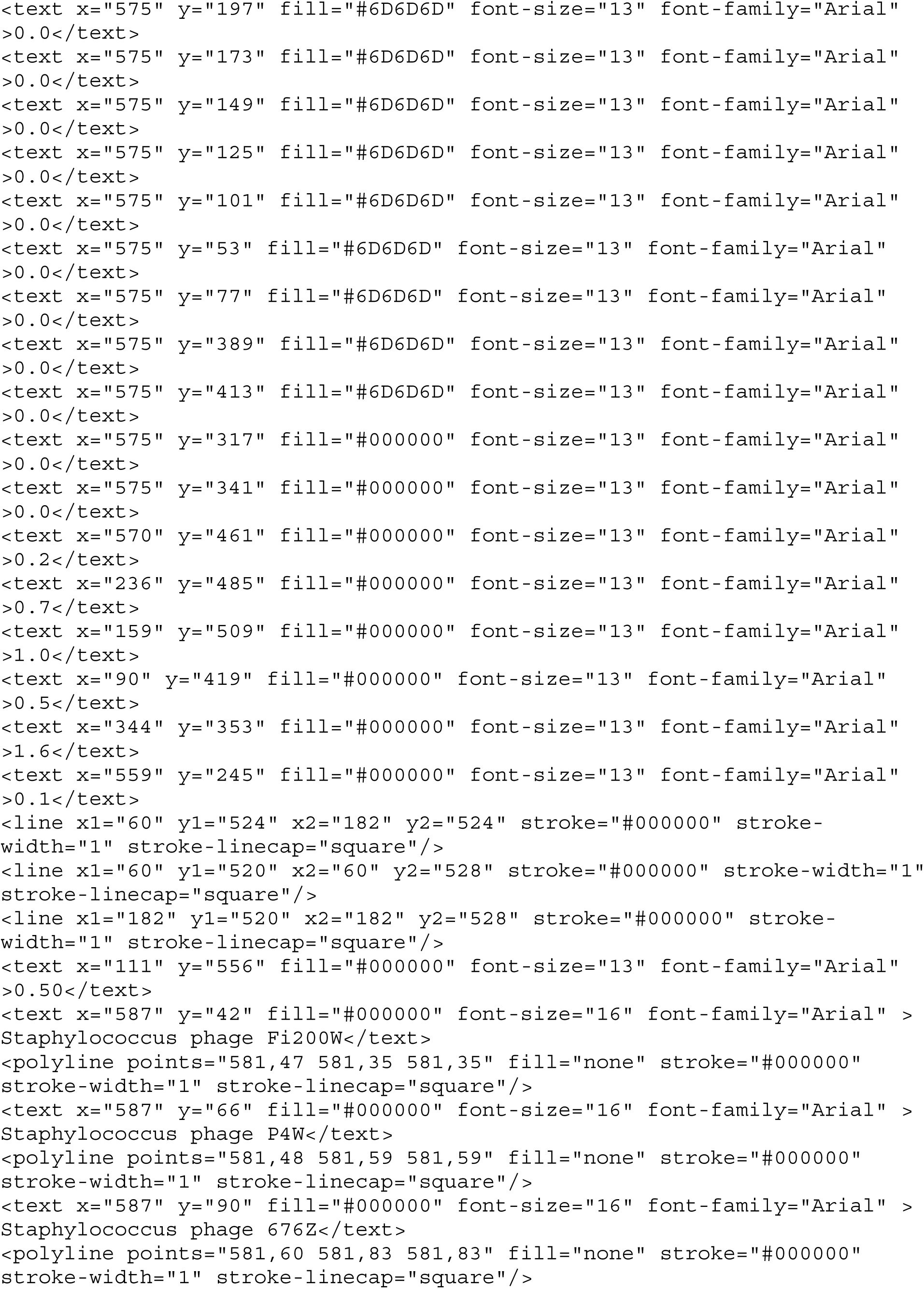
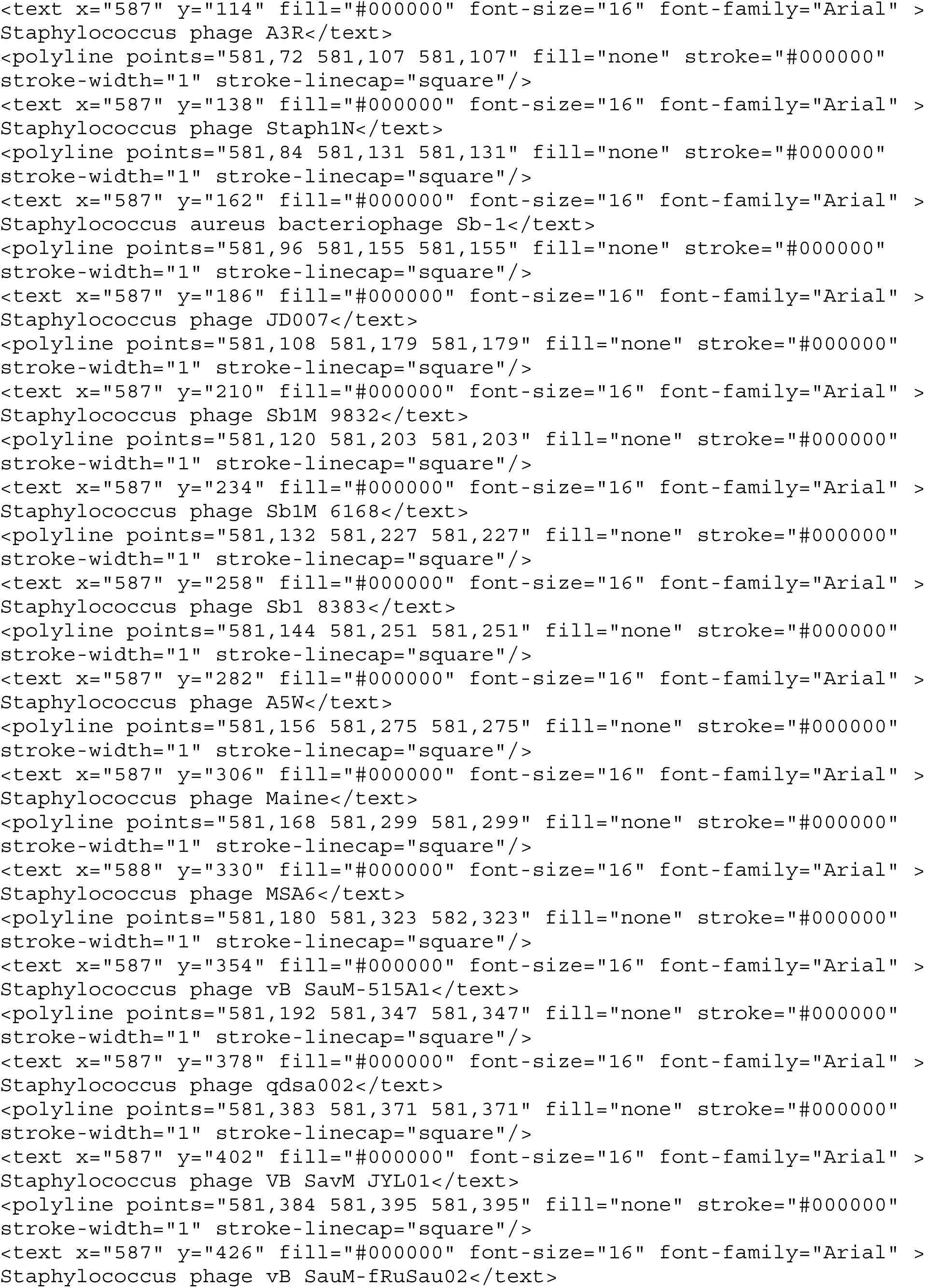
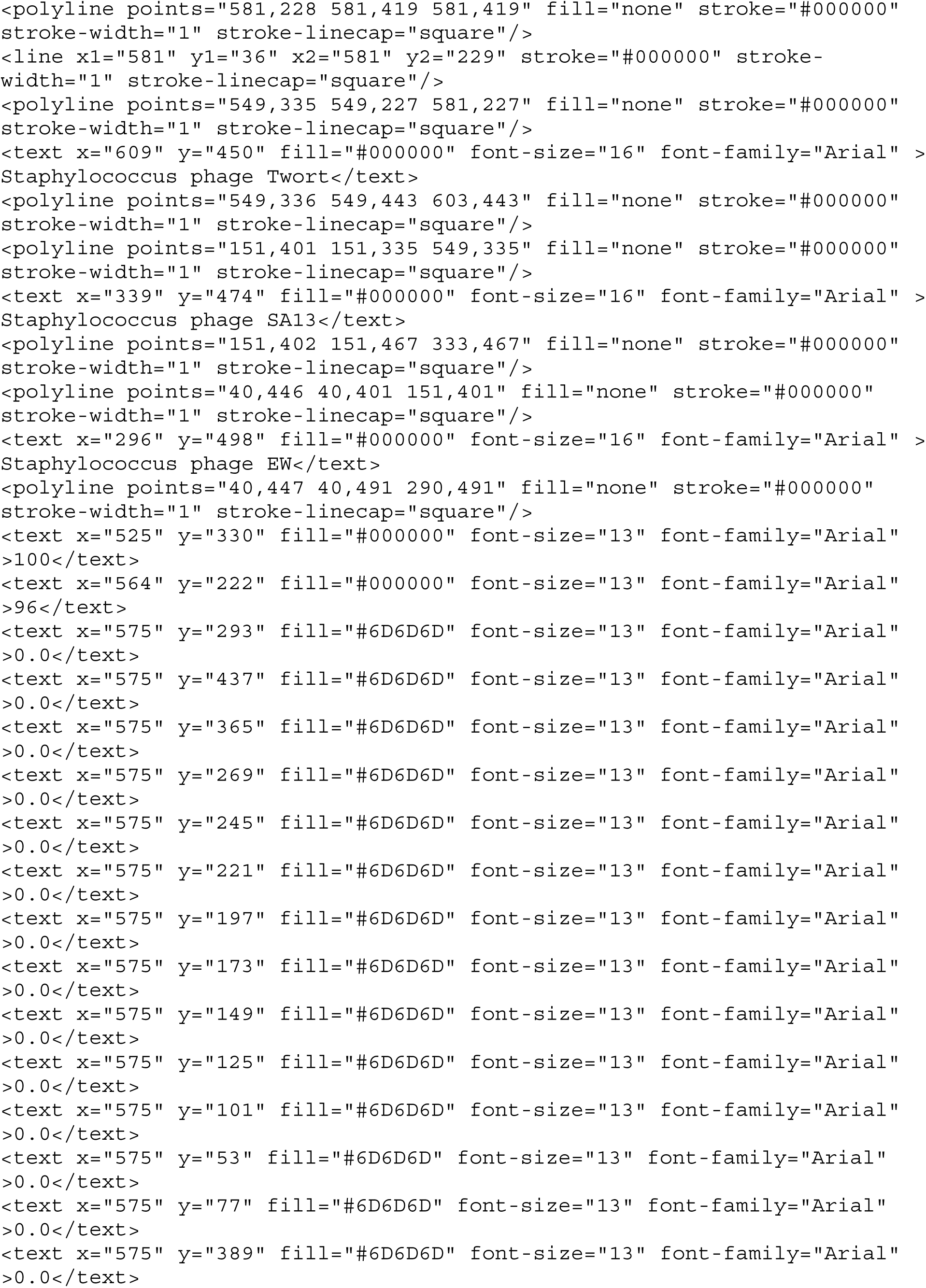
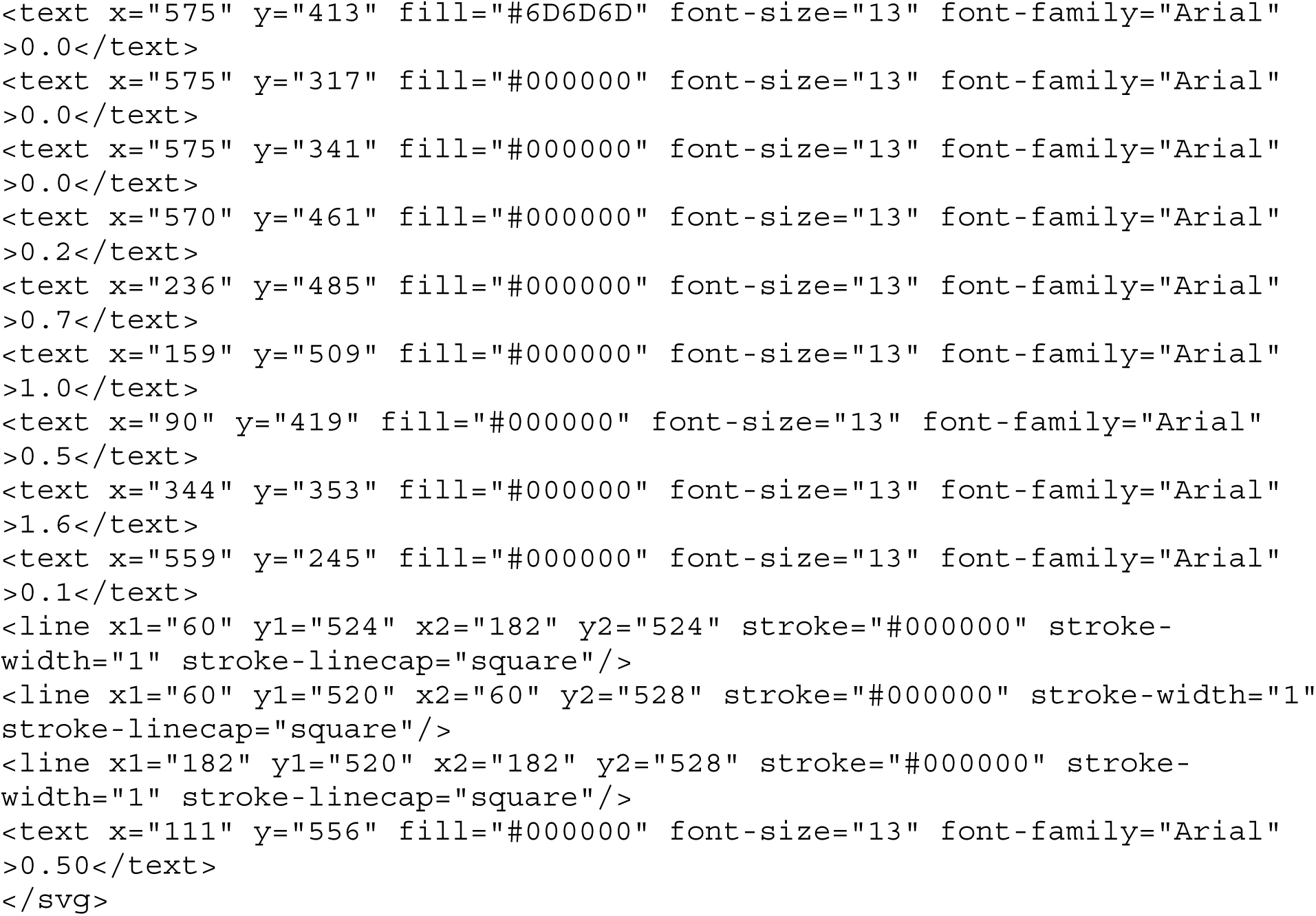

## Supporting information

https://github.com/HivanA98/In-silico-characterization-of-lysis-and-host-recognition-modules-in-S.-aureus-bacteriophage-genomes

## Footnotes

1 NCBI (2017). GenBank Overview [online]. Website https://www.ncbi.nlm.nih.gov/genbank/ [accessed 30 May 2026].

