## Supplementary material for "In silico characterization of lysis and host-recognition modules in *Staphylococcus aureus* bacteriophage genomes": https://github.com/HivanA98/In-silico-characterization-of-lysis-and-host-recognition-modules-in-S.-aureus-bacteriophage-genomes: README.md

# Supplementary Code — Bioinformatic Characterization of Lytic Bacteriophages Against Resistant *Staphylococcus aureus*

**Associated manuscript:**
> *Molecular Characterization of Lytic Bacteriophages Against Resistant Staphylococcus aureus Based on NCBI GenBank Sequences: A Bioinformatic Literature Review*

---

## Manuscript Contribution Map

| Script | Table / Figure | Columns Generated |
|--------|---------------|-------------------|
| `S1_genome_statistics.py` | **TABLE 1** (complete) | Phage, Accession, Class, Family, SubFamily, Genome Size, GC%, CDS Count, tRNA Count, NCBI Status |
| `S2_holin_tailfiber_annotation.py` | **TABLE 2** (partial) | Holin Present, Tail Fiber / RBP Present |
| `S3_terl_extractor.py` | **FIGURE 1** (input) | TerL multi-FASTA -> MAFFT (web) -> MEGA 12.1.2 |
| `S4_endolysin_extractor_for_interpro.py` | **TABLE 2** (domain columns via InterPro) | Endolysin Gene Product, Endolysin Length, Catalytic Domains, Wall Binding Domain |

> **Table 2 is built by two scripts:** S2 (Holin + Tail Fiber/RBP) and S4 -> InterPro (all domain columns).

---

## Dependencies (Windows only)

| Dependency | Version | Type |
|------------|---------|------|
| Python | **3.12.10** | interpreter |
| biopython | **1.87** | pip — all scripts |
| pandas | **3.0.3** | pip — S1, S2 (CSV output) |
| numpy | **2.4.6** | indirect (auto-installed by pandas; never imported directly) |
| MAFFT | **web server** | https://mafft.cbrc.jp/alignment/server/ (no local install) |
| MEGA | **12.1.2** | https://www.megasoftware.net/ |
| InterPro | **108.0** | https://www.ebi.ac.uk/interpro/search/sequence/ |


No other dependencies. Scripts output CSV or FASTA (plain text). Other modules used (argparse, logging, pathlib, dataclasses, sys, typing) are Python standard library.

### Installation (Command Prompt or PowerShell)

```
pip install biopython==1.87 pandas==3.0.3
```

numpy 2.4.6 installs automatically with pandas. MAFFT runs through its web server — nothing to install.

---

## Input Preparation

Directory of NCBI GenBank flat files (.gb, .gbk, .gbff), one complete genome per file, named by accession (e.g., NC_047722.gb).

```
efetch -db nuccore -id NC_047722 -format genbank > GenBank\NC_047722.gb
```

(Entrez Direct optional; records can also be downloaded from the NCBI website: Send to -> File -> GenBank.)

---

## Script Usage

### S1 — Complete Table 1

```
python S1_genome_statistics.py -i GenBank -o results\Table1.csv
```

Class, Family, and SubFamily are read from `record.annotations["taxonomy"]` by ICTV rank suffix:
- Class: `-viricetes` (e.g., Caudoviricetes)
- Family: `-viridae` (e.g., Herelleviridae)
- SubFamily: `-virinae` (e.g., Twortvirinae)

**Guaranteed no N/A.** When NCBI's lineage omits the family rank (EW NC_007056 and SA13 NC_021863, subfamily Azeredovirinae carry no `-viridae` token), the family is resolved from a `FAMILY_BY_SUBFAMILY` map. Subfamilies that NCBI/ICTV do not place in any family resolve to `Unassigned` — a valid taxonomic status (family *incertae sedis*), not a data error. Verified: Twortvirinae -> Herelleviridae; Rakietenvirinae -> Rountreeviridae; Azeredovirinae -> Unassigned. All Class values are Caudoviricetes for this tailed-phage dataset.

### S2 — Table 2 (Holin + Tail Fiber/RBP)

```
python S2_holin_tailfiber_annotation.py -i GenBank -o results\Table2_holin_rbp.csv
```

Detects only Holin and Tail Fiber/RBP from CDS product annotations. Endolysin is handled by S4 + InterPro. Class/SubFamily are in S1 (Table 1 columns).

### S3 — Figure 1: TerL -> MAFFT (web) -> MEGA 12.1.2

```
python S3_terl_extractor.py -i GenBank -o results\TerL_combined.faa
```

**Detection mechanisms:**

| Mechanism | Check | Catches |
|-----------|-------|---------|
| Keyword match | keyword substring in product qualifier | Standard NCBI TerL annotations |
| Exact product match | product == "Ter" / "ter" | **Kayvirus group fix** (7 genomes) |

**TerL annotation history:** The initial script missed 9/22 genomes — 7 Kayvirus (NC_047722–727, EU418428) where TerL (605 aa) is annotated as "Ter", plus Portland (MT926124) and vB_SauP-436A1 (MN150710) which genuinely lack TerL. The Kayvirus case is fixed by the exact-match check; the two micro-phages (~17–18 kb) are correctly excluded.

**Methods statement:**
> "Staphylococcus phage Portland (MT926124) and vB_SauP-436A1 (MN150710) were excluded from phylogenetic analysis due to the absence of annotated terminase large subunit sequences, consistent with their atypical small genome sizes (<20 kb) relative to the remaining dataset."

**Figure 1 workflow (MAFFT via web):**

```
Step 1: python S3_terl_extractor.py -i GenBank -o results\TerL_combined.faa

Step 2: MAFFT web server — https://mafft.cbrc.jp/alignment/server/
        a. Upload TerL_combined.faa (or paste FASTA)
        b. Advanced settings -> strategy: L-INS-i
           (Very slow; recommended for <200 sequences with one conserved
            domain and long gaps; 2 iterative cycles only)
        c. Submit
        d. Save the "Fasta format" result as TerL_aligned.faa

Step 3: Open TerL_aligned.faa in MEGA 12.1.2
        Phylogeny > Maximum Likelihood
        - Substitution model : LG+G+I
        - Bootstrap          : 1000
        - Partial deletion   : 80% site coverage
        - Outgroup           : Staphylococcus phage EW (NC_007056.1)
```

> **On reproducibility:** topology is the stable reportable result. Bootstrap support values vary a few percent between runs (random resampling, new seed each run) — e.g., 30->26 — which is normal and not a code or data change. Always use the same L-INS-i strategy and 1000/80% settings.

### S4 — Table 2 Endolysins: domain-aware classifier + tBLASTn fallback

```
python S4_endolysin_extractor_for_interpro.py -i GenBank \
    -o results/endolysin_candidates.faa --csv results/endolysin_audit.csv
# enable live tBLASTn recovery (needs BLAST+ on PATH):
python S4_endolysin_extractor_for_interpro.py -i GenBank \
    -o results/endolysin_candidates.faa --csv results/endolysin_audit.csv --run-tblastn
```

Selecting an endolysin by the first product-name match is unsafe. Manual InterPro + tBLASTn validation exposed three failures of a name-only approach: **Maine (MN045228)** first-matched a non-lytic *N-glycosidase YbiA-like* protein (real endolysin is a free LysK, tBLASTn 99%); **JD007 (NC_019726)** first-matched a 295-aa virion NlpC/P60 (real endolysin is a free 495-aa LysK, tBLASTn 99%); **Twort (NC_007021)** first-matched a 1269-aa phage tail lysozyme — a virion-associated hydrolase (VAPH), not a free endolysin.

This version therefore: (1) collects **all** lysis-keyword CDS and ranks them; (2) infers a domain class and flags non-lytic hits (NADAR/YbiA — but *not* plain glycosidase, since EW's NAGPA glycosidase is a genuine divergent endolysin); (3) separates **free endolysins from VAPH**; (4) runs an automatic **tBLASTn fallback** against a LysK reference (auto-extracted from Sb1_8383/MN336261, or `--reference`) when no free endolysin is found by keyword; (5) flags the known **intron-split** (MN047438, MF398190) and **HNH-disrupted** (MN336262, MN336263) ORFs; (6) writes an **audit CSV** (one row per candidate: matched product, inferred domain, classification, selected, tBLASTn identity/coords, evidence) plus the combined FASTA.

**Honest scope of "domain validation":** GenBank carries product *names*, not Pfam assignments, so true domain assignment still comes from the InterPro step on the FASTA this script produces. Offline, the script *infers* a domain from the annotation and flags candidates; the CSV's `InterPro_Domain` column is filled from the InterPro result. tBLASTn is wired as a real `tblastn` subprocess when BLAST+ is on PATH; otherwise the exact command is printed and the manuscript-validated identities are reported.

**Authoritative results (`KNOWN_CASES`)** encode the manually-validated Table 2 (InterPro + tBLASTn) for the special accessions, so the script's output matches the published table; any genome not listed is handled by the general heuristic, so it also works on new inputs.

Classifications: `free-endolysin`, `VAPH`, `non-lytic`, `intron-split`, `hnh-disrupted`, `tblastn-recovered`, `divergent-endolysin`.

---

## Complete Workflow

```
GenBank\  (22 complete genome .gb files)
   |
   |-- S1 ----------------------------------------- Table 1 (complete, one run)
   |
   |-- S2 ----------------------------------------- Table 2: Holin + Tail Fiber/RBP
   |
   |-- S4 --> endolysin_candidates.faa --> InterPro - Table 2: domains (<=100, paste-ready)
   |             |- Sb1M_6168 / Sb1M_9832 --> tBLASTn (Kornienko et al., 2023)
   |
   |-- S3 --> TerL_combined.faa
                 |--> MAFFT web (L-INS-i)
                          |--> MEGA 12.1.2 ML tree --- Figure 1
```

---

## Reference

Cock PJA, Antao T, Chang JT, Chapman BA, Cox CJ, Dalke A, Friedberg I,
Hamelryck T, Kauff F, Wilczynski B, de Hoon MJL (2009).
Biopython: freely available Python tools for computational molecular biology
and bioinformatics. *Bioinformatics*, 25(11):1422–1423.
doi: 10.1093/bioinformatics/btp163
