## Supplementary material for "In silico characterization of lysis and host-recognition modules in *Staphylococcus aureus* bacteriophage genomes": https://github.com/HivanA98/In-silico-characterization-of-lysis-and-host-recognition-modules-in-S.-aureus-bacteriophage-genomes: S1_genome_statistics.py

```
#!/usr/bin/env python3
"""
S1_genome_statistics.py
=======================
Batch Genomic Statistics Extractor — Complete Table 1 Generator

==============================================================================
MANUSCRIPT CONTRIBUTION
  Generates:  TABLE 1 — General Characteristics of Phage Genomes
  Output columns (all):
    Phage | Accession | Class | SubFamily | Genome_Size_bp | GC_Percent |
    CDS_Count | tRNA_Count | NCBI_Status
==============================================================================

Associated manuscript:
    "Molecular Characterization of Lytic Bacteriophages Against Resistant
    Staphylococcus aureus Based on NCBI GenBank Sequences:
    A Bioinformatic Literature Review"

Description
-----------
Extracts all Table 1 data from GenBank flat files in a single run:

  1. Genomic statistics (genome size, GC%, CDS count, tRNA count):
     Derived from record.seq (nucleotide sequence) and record.features
     (list of SeqFeature objects), following Cock et al. (2009).

     CDS COUNT — METHOD NOTE (reconciliation with primary literature):
     CDS_Count is the number of features with feature.type == "CDS" in the
     GenBank record AS DEPOSITED. It is therefore the curated annotation the
     submitter uploaded, read verbatim by Biopython SeqIO — NOT a re-prediction.
     Primary papers that re-annotate a genome with a de-novo gene-caller
     (e.g., RAST, Prokka, GeneMarkS) can report a DIFFERENT ORF count for the
     same accession. Example in this dataset: Staphylococcus phage vB_SauM-515A1
     (MN047438) is reported with 238 ORFs by RAST in Kornienko et al. (2020,
     Sci Rep and Viruses), whereas the deposited GenBank record yields 236 CDS.
     The 2-feature difference reflects gene-caller and curation differences
     (and is compounded for MN047438 by its intron-split lysK; see S4) — it is
     NOT a counting error in this script. Matching 238 would require re-running
     RAST, which is a different methodology from "characterization of the
     deposited GenBank annotation". Report CDS as the GenBank-deposit count and
     add a manuscript footnote stating the method (see README / docx).

  2. Taxonomic classification (Class, Family, SubFamily):
     Derived from record.annotations["taxonomy"], the LINEAGE field in
     NCBI GenBank records, using ICTV rank suffixes:
       Class     ← token ending in "-viricetes"  (e.g., Caudoviricetes)
       Family    ← token ending in "-viridae"    (e.g., Herelleviridae)
       SubFamily ← token ending in "-virinae"     (e.g., Twortvirinae)

     This corrects an earlier version that mislabelled the "-viridae"
     family rank as "Class". Under ICTV nomenclature the hierarchy is:
       Caudoviricetes  (-viricetes)  CLASS
       Caudovirales    (-virales)    ORDER  (abolished in ICTV 2022 for
                                             many tailed-phage lineages)
       Herelleviridae  (-viridae)    FAMILY
       Twortvirinae    (-virinae)    SUBFAMILY
       Kayvirus        (-virus)      GENUS

     Guaranteed no "N/A": some NCBI lineages omit the family rank
     (e.g., subfamily Azeredovirinae for phages EW [NC_007056] and
     SA13 [NC_021863] carries no "-viridae" token). For these records the
     family is filled from FAMILY_BY_SUBFAMILY; subfamilies that NCBI/ICTV
     do not place in any family resolve to "Unassigned" — a valid taxonomic
     status (family incertae sedis), NOT a data error. Edit
     FAMILY_BY_SUBFAMILY if ICTV later assigns a family.

     Verified mapping (this dataset):
       Twortvirinae    → Family Herelleviridae,  Class Caudoviricetes
       Rakietenvirinae → Family Rountreeviridae,  Class Caudoviricetes
       Azeredovirinae  → Family Unassigned,       Class Caudoviricetes

  3. Completeness status (NCBI_Status):
     Inferred from KEYWORDS and DEFINITION fields.

Reference
---------
Cock PJA, Antao T, Chang JT, Chapman BA, Cox CJ, Dalke A, Friedberg I,
Hamelryck T, Kauff F, Wilczynski B, de Hoon MJL (2009). Biopython: freely
available Python tools for computational molecular biology and bioinformatics.
Bioinformatics, 25(11):1422-1423. doi:10.1093/bioinformatics/btp163

Dependencies (exact versions; no other dependencies required)
--------------------------------------------------------------
  Installed via pip:
    biopython  1.87    GenBank parsing (Cock et al., 2009)
    pandas     3.0.3   tabular CSV output
    numpy      2.4.6   INDIRECT — installed automatically by pandas;
                       NOT imported directly by this script.

  Python standard library (no installation needed):
    argparse, logging, sys, pathlib, typing

  Output is CSV only; the dependency set is exactly the components above.

Tested Environment (Windows)
-----------------------------
    OS        : Windows 10 / 11
    Python    : 3.12.10
    biopython : 1.87
    pandas    : 3.0.3
    numpy     : 2.4.6  (indirect, via pandas)

Installation (Command Prompt / PowerShell)
------------------------------------------
    pip install biopython==1.87 pandas==3.0.3
    (numpy 2.4.6 is installed automatically as a pandas dependency)

Usage (Windows Command Prompt)
-------------------------------
    python S1_genome_statistics.py -i GenBank -o results/Table1.csv
"""

import argparse
import logging
import sys
from pathlib import Path
from typing import Optional

import pandas as pd
from Bio import SeqIO
from Bio.SeqRecord import SeqRecord

# ---------------------------------------------------------------------------
# Logging
# ---------------------------------------------------------------------------
logging.basicConfig(
    level=logging.INFO,
    format="%(asctime)s  [%(levelname)-8s]  %(message)s",
    datefmt="%Y-%m-%d %H:%M:%S",
)
log = logging.getLogger(__name__)

# ---------------------------------------------------------------------------
# Constants
# ---------------------------------------------------------------------------
VALID_EXTENSIONS: frozenset[str] = frozenset({".gb", ".gbk", ".gbff"})

# Fallback family lookup for records whose NCBI lineage OMITS the family
# ("-viridae") rank. Some NCBI lineages list only down to subfamily; in that
# case the family is resolved from the subfamily below.
#
# Values reflect ICTV placement as retrieved from the NCBI Taxonomy browser:
#   - Twortvirinae   is placed in family Herelleviridae.
#   - Rakietenvirinae is placed in family Rountreeviridae.
#   - Azeredovirinae  is NOT placed in any family in the NCBI lineage for
#     phages EW (NC_007056) and SA13 (NC_021863); its family-level rank is
#     unresolved (incertae sedis), so it resolves to "Unassigned".
#
# "Unassigned" is a valid taxonomic status, NOT a missing-data marker.
# If ICTV later assigns a family to a subfamily here, edit this map.
FAMILY_BY_SUBFAMILY: dict[str, str] = {
    "Twortvirinae":    "Herelleviridae",
    "Rakietenvirinae": "Rountreeviridae",
    "Azeredovirinae":  "Unassigned",
}

# Default class for tailed dsDNA bacteriophages (used only if a lineage
# unexpectedly lacks a "-viricetes" token; every genome in this dataset
# carries Caudoviricetes in its lineage).
DEFAULT_CLASS: str = "Caudoviricetes"


# ===========================================================================
# Core analysis functions
# ===========================================================================

def calculate_gc_content(sequence: str) -> float:
    """
    Calculate percentage GC content of a nucleotide sequence.

    Formula: GC% = (count_G + count_C) / len(sequence) × 100

    Ambiguity codes (N, R, Y, etc.) count toward the denominator but not
    the numerator, consistent with NCBI GenBank composition reporting.

    Parameters
    ----------
    sequence : str
        Raw nucleotide sequence (any case).

    Returns
    -------
    float
        GC percentage rounded to two decimal places. Returns 0.00 if empty.
    """
    seq   = sequence.upper()
    total = len(seq)
    if total == 0:
        return 0.00
    gc = seq.count("G") + seq.count("C")
    return round((gc / total) * 100, 2)


def extract_classification(record: SeqRecord) -> tuple[str, str, str]:
    """
    Extract ICTV Class, Family, and SubFamily from a SeqRecord.

    The NCBI GenBank LINEAGE field is mapped to record.annotations["taxonomy"]
    by Biopython as a list of taxonomic strings in hierarchical order.

    Rank detection (by suffix):
      Class     ← token ending in "-viricetes"  (e.g., Caudoviricetes)
      Family    ← token ending in "-viridae"    (e.g., Herelleviridae)
      SubFamily ← token ending in "-virinae"     (e.g., Twortvirinae)

    No-"N/A" guarantee:
      - If no "-viricetes" token is present, Class falls back to
        DEFAULT_CLASS ("Caudoviricetes"); every genome in this dataset
        carries Caudoviricetes, so this fallback is a safety net only.
      - If no "-viridae" token is present (NCBI omits the family rank, as for
        the Azeredovirinae phages EW and SA13), Family is resolved from
        FAMILY_BY_SUBFAMILY; unresolved subfamilies yield "Unassigned".
      - If no "-virinae" token is present, SubFamily yields "Unassigned".

    Parameters
    ----------
    record : SeqRecord
        A Biopython SeqRecord parsed from a GenBank flat file.

    Returns
    -------
    tuple[str, str, str]
        (Class, Family, SubFamily). No element is ever the literal "N/A".

    Notes
    -----
    Representative lineages in this dataset:
      Caudoviricetes; Herelleviridae;  Twortvirinae    → Kayvirus / Twortvirus
      Caudoviricetes; Rountreeviridae; Rakietenvirinae → Andhravirus
      Caudoviricetes; <no family>;     Azeredovirinae  → (family Unassigned)
    """
    taxonomy = record.annotations.get("taxonomy", [])

    phage_class = None
    family      = None
    subfamily   = None

    for taxon in taxonomy:
        if taxon.endswith("viricetes"):
            phage_class = taxon
        elif taxon.endswith("viridae"):
            family = taxon
        elif taxon.endswith("virinae"):
            subfamily = taxon

    # ---- Resolve with no-"N/A" fallbacks ----
    if phage_class is None:
        phage_class = DEFAULT_CLASS

    if subfamily is None:
        subfamily = "Unassigned"

    if family is None:
        # NCBI omitted the family rank: resolve from the subfamily map.
        family = FAMILY_BY_SUBFAMILY.get(subfamily, "Unassigned")

    return phage_class, family, subfamily


def infer_ncbi_status(record: SeqRecord) -> str:
    """
    Infer NCBI sequence completeness from KEYWORDS and DEFINITION fields.

    Returns
    -------
    str
        "Complete Genome" if "complete" appears in keywords or description;
        "Draft/Partial" otherwise.
    """
    keywords    = record.annotations.get("keywords", [])
    description = record.description.lower()

    if any("complete" in kw.lower() for kw in keywords) or "complete" in description:
        return "Complete Genome"
    return "Draft/Partial"


def extract_stats(record: SeqRecord) -> dict:
    """
    Extract all Table 1 fields from a single SeqRecord.

    Parameters
    ----------
    record : SeqRecord
        A Biopython SeqRecord from Bio.SeqIO.read().

    Returns
    -------
    dict
        Keys: Phage, Accession, Class, Family, SubFamily, Genome_Size_bp,
        GC_Percent, CDS_Count, tRNA_Count, NCBI_Status.

    Notes
    -----
    CDS and tRNA counts iterate over record.features and filter by
    feature.type. Only "CDS" and "tRNA" type annotations are counted;
    other feature types (gene, repeat_region, misc_feature) are ignored.
    """
    seq_str      = str(record.seq).upper()
    cds_count    = sum(1 for f in record.features if f.type == "CDS")
    trna_count   = sum(1 for f in record.features if f.type == "tRNA")
    cls, family, subfam = extract_classification(record)

    return {
        "Phage":          record.annotations.get("organism", record.name),
        "Accession":      record.id,
        "Class":          cls,
        "Family":         family,
        "SubFamily":      subfam,
        "Genome_Size_bp": len(seq_str),
        "GC_Percent":     calculate_gc_content(seq_str),
        "CDS_Count":      cds_count,
        "tRNA_Count":     trna_count,
        "NCBI_Status":    infer_ncbi_status(record),
    }


def parse_single_file(gb_path: Path) -> Optional[dict]:
    """
    Parse one GenBank file and return Table 1 row data.

    Bio.SeqIO.read() is used (not parse()) because each file must contain
    exactly one complete genome record.

    Parameters
    ----------
    gb_path : Path
        Path to a GenBank flat file.

    Returns
    -------
    dict or None
        Row data on success; None if parsing fails. Failures are logged at
        WARNING level and the file is skipped without halting execution.
    """
    try:
        record = SeqIO.read(str(gb_path), "genbank")
        return extract_stats(record)
    except ValueError as exc:
        log.warning(f"  Skipped '{gb_path.name}': {exc}")
    except Exception as exc:
        log.warning(f"  Skipped '{gb_path.name}': unexpected error — {exc}")
    return None


# ===========================================================================
# Batch processing and I/O
# ===========================================================================

def run(input_dir: Path, output_path: Path) -> pd.DataFrame:
    """
    Process all GenBank files and write the complete Table 1.

    Parameters
    ----------
    input_dir : Path
        Directory containing GenBank flat files (.gb / .gbk / .gbff).
        Sub-directories are not traversed.
    output_path : Path
        Output CSV file path (UTF-8 encoded).

    Returns
    -------
    pd.DataFrame
        Complete Table 1 sorted by Accession number.
    """
    gb_files = sorted(
        f for f in input_dir.iterdir()
        if f.is_file() and f.suffix.lower() in VALID_EXTENSIONS
    )

    if not gb_files:
        log.error(
            f"No GenBank files found in '{input_dir}'. "
            f"Accepted: {', '.join(sorted(VALID_EXTENSIONS))}"
        )
        sys.exit(1)

    log.info(f"Found {len(gb_files)} GenBank file(s) in '{input_dir}'")

    rows = []
    for gb in gb_files:
        log.info(f"  Processing {gb.name}")
        row = parse_single_file(gb)
        if row:
            rows.append(row)

    if not rows:
        log.error("No records extracted. Verify input files are valid GenBank format.")
        sys.exit(1)

    df = pd.DataFrame(rows).sort_values("Accession").reset_index(drop=True)

    output_path.parent.mkdir(parents=True, exist_ok=True)
    df.to_csv(output_path, index=False, encoding="utf-8")

    log.info(f"Table 1 written: '{output_path}'  ({len(df)} records)")
    return df


# ===========================================================================
# Command-line interface
# ===========================================================================

def build_parser() -> argparse.ArgumentParser:
    parser = argparse.ArgumentParser(
        prog="S1_genome_statistics.py",
        description=(
            "Extract all Table 1 data from Staphylococcus phage GenBank records: "
            "taxonomic classification (Class, Family, SubFamily) and genomic statistics "
            "(genome size, GC%%, CDS count, tRNA count). "
            "This script generates the COMPLETE Table 1 in a single run."
        ),
        formatter_class=argparse.RawDescriptionHelpFormatter,
        epilog="""
MANUSCRIPT OUTPUT
-----------------
  TABLE 1 — General Characteristics of Phage Genomes
  Columns: Phage | Accession | Class | SubFamily | Genome Size | GC%% |
           CDS Count | tRNA Count | NCBI Status

Examples (Windows Command Prompt)
----------------------------------
  python S1_genome_statistics.py -i GenBank -o results\\Table1.csv

Examples (Windows PowerShell)
------------------------------
  python S1_genome_statistics.py -i GenBank -o results/Table1.csv
        """,
    )
    parser.add_argument(
        "--input_dir", "-i",
        type=Path, required=True, metavar="DIR",
        help="Directory containing GenBank files (.gb / .gbk / .gbff)",
    )
    parser.add_argument(
        "--output", "-o",
        type=Path, default=Path("Table1.csv"), metavar="FILE",
        help="Output CSV file path. Default: Table1.csv",
    )
    return parser


# ===========================================================================
# Entry point
# ===========================================================================

if __name__ == "__main__":
    args = build_parser().parse_args()
    df   = run(args.input_dir, args.output)

    sep = "=" * 80
    print(f"\n{sep}")
    print("TABLE 1 — GENERAL CHARACTERISTICS OF PHAGE GENOMES")
    print(sep)
    print(df.to_string(index=False))
    print(sep)
    print(f"  Total phages       : {len(df)}")
    print(f"  Genome size range  : {df['Genome_Size_bp'].min():,} – "
          f"{df['Genome_Size_bp'].max():,} bp")
    print(f"  GC%% range          : {df['GC_Percent'].min():.2f}%% – "
          f"{df['GC_Percent'].max():.2f}%%")
    print(f"  Classes found      : "
          f"{', '.join(sorted(df['Class'].unique()))}")
    print(f"  Families found     : "
          f"{', '.join(sorted(df['Family'].unique()))}")
    print(f"  SubFamilies found  : "
          f"{', '.join(sorted(df['SubFamily'].unique()))}")
    # Verify no residual N/A in any taxonomic column
    na_mask = (df[["Class", "Family", "SubFamily"]] == "N/A").any(axis=1)
    print(f"  Rows with 'N/A'    : {int(na_mask.sum())}  (target: 0)")
    print(sep)
```
