## Supplementary material for "In silico characterization of lysis and host-recognition modules in *Staphylococcus aureus* bacteriophage genomes": https://github.com/HivanA98/In-silico-characterization-of-lysis-and-host-recognition-modules-in-S.-aureus-bacteriophage-genomes: S2_holin_tailfiber_annotation.py

```
#!/usr/bin/env python3
"""
S2_holin_tailfiber_annotation.py
=================================
Holin and Tail Fiber / Receptor-Binding Protein Annotator

==============================================================================
MANUSCRIPT CONTRIBUTION
  Generates partial data for:
    TABLE 2 — Conservation of Lysis Genes and Variability of
              Host-Recognition Modules
  Output columns used from this script:
    Holin Present | Tail Fiber / RBP Present

  The remaining Table 2 columns (Endolysin Gene Product, Endolysin Length,
  Catalytic Domains, Wall Binding Domain) are produced by S4_endolysin_extractor_for_interpro.py
  followed by InterPro domain annotation. See S4 documentation.
==============================================================================

Associated manuscript:
    "Molecular Characterization of Lytic Bacteriophages Against Resistant
    Staphylococcus aureus Based on NCBI GenBank Sequences:
    A Bioinformatic Literature Review"

Description
-----------
Scans CDS product annotations in each GenBank record to detect two functional
gene categories critical to bacteriophage lytic activity:

  1. Holin — membrane-perforating protein that gates endolysin release into
     the periplasm during the lytic cycle. Detected by keyword matching in
     the "product" qualifier of CDS SeqFeature objects.

  2. Tail fiber / Receptor-Binding Protein (RBP) — host-recognition protein
     located at the phage tail that determines host specificity by binding
     specific surface receptors. Variability in tail fiber/RBP genes directly
     explains differences in host range among phages.

This script does NOT perform endolysin detection. Endolysin identification
and domain architecture (CHAP, Amidase, SH3b, LysM) are handled by
S4_endolysin_extractor_for_interpro.py followed by InterPro analysis.

Detection Method
----------------
Detection is keyword-based, matching substrings (case-insensitive) in the
concatenated text of the "product", "gene", "note", and "function" qualifiers
of each CDS SeqFeature (Cock et al., 2009).

Limitation
----------
This script detects only what is explicitly annotated as text in GenBank
flat files. Unannotated or hypothetically annotated holin or tail fiber
sequences will not be detected. Absence of detection ("No") does not
definitively indicate biological absence; it may reflect incomplete
annotation in the GenBank record.

Reference
---------
Cock PJA, Antao T, Chang JT, Chapman BA, Cox CJ, Dalke A, Friedberg I,
Hamelryck T, Kauff F, Wilczynski B, de Hoon MJL (2009). Biopython: freely
available Python tools for computational molecular biology and bioinformatics.
Bioinformatics, 25(11):1422-1423. doi:10.1093/bioinformatics/btp163

Dependencies (exact versions; no other dependencies required)
--------------------------------------------------------------
  Installed via pip:
    biopython  1.87    GenBank parsing (Cock et al., 2009)
    pandas     3.0.3   tabular CSV output
    numpy      2.4.6   INDIRECT — installed automatically by pandas;
                       NOT imported directly by this script.

  Python standard library (no installation needed):
    argparse, logging, sys, pathlib, typing

  Output is CSV only.

Tested Environment (Windows)
-----------------------------
    OS        : Windows 10 / 11
    Python    : 3.12.10
    biopython : 1.87
    pandas    : 3.0.3
    numpy     : 2.4.6  (indirect, via pandas)

Installation (Command Prompt / PowerShell)
------------------------------------------
    pip install biopython==1.87 pandas==3.0.3
    (numpy 2.4.6 is installed automatically as a pandas dependency)

Usage (Windows Command Prompt)
-------------------------------
    python S2_holin_tailfiber_annotation.py -i GenBank -o results/Table2_holin_rbp.csv
"""

import argparse
import logging
import sys
from pathlib import Path
from typing import Optional

import pandas as pd
from Bio import SeqIO
from Bio.SeqRecord import SeqRecord

# ---------------------------------------------------------------------------
# Logging
# ---------------------------------------------------------------------------
logging.basicConfig(
    level=logging.INFO,
    format="%(asctime)s  [%(levelname)-8s]  %(message)s",
    datefmt="%Y-%m-%d %H:%M:%S",
)
log = logging.getLogger(__name__)

# ---------------------------------------------------------------------------
# Constants
# ---------------------------------------------------------------------------
VALID_EXTENSIONS: frozenset[str] = frozenset({".gb", ".gbk", ".gbff"})

# Holin detection keywords
# Matched as case-insensitive substrings in CDS qualifier text.
HOLIN_KEYWORDS: tuple[str, ...] = (
    "holin",
    "putative holin",
    "class i holin",
    "class ii holin",
    "phage holin",
)

# Tail fiber / Receptor-Binding Protein (RBP) detection keywords
# RBP variability directly determines host range and is the primary target
# for synthetic biology engineering in phage therapy development.
#
# IMPORTANT — "tail tube protein" is deliberately EXCLUDED. The tail tube is a
# structural conduit through which DNA is ejected; it is present in essentially
# all tailed phages and is NOT a receptor-binding / host-recognition protein.
# An earlier draft of this script included "tail tube protein", which produced
# false positives (5 genomes incorrectly flagged Yes vs the pre-refactor code:
# MN336261, MN336262, MN336263, NC_047725, NC_047726). It has been removed.
RBP_KEYWORDS: tuple[str, ...] = (
    "tail fiber",
    "tail fibre",
    "receptor binding protein",
    "receptor-binding protein",
    "rbp",
    "tail spike",
    "tailspike",
    "tail-spike",
    "host recognition protein",
    "adsorption protein",
    "baseplate receptor-binding",
    "host specificity protein",
)


# ===========================================================================
# Helper utilities
# ===========================================================================

def get_qualifier_text(feature) -> str:
    """
    Concatenate CDS qualifier values into one searchable string.

    Qualifiers searched: product, gene, note, function.
    Returned text is lowercased for case-insensitive matching.

    Parameters
    ----------
    feature : SeqFeature
        A Biopython SeqFeature object (type "CDS").

    Returns
    -------
    str
        Lowercased, space-joined qualifier values.
    """
    parts = []
    for key in ("product", "gene", "note", "function"):
        parts.extend(feature.qualifiers.get(key, []))
    return " ".join(parts).lower()


# ===========================================================================
# Per-genome detection
# ===========================================================================

def _match_keyword(feature, keywords: tuple[str, ...]) -> Optional[tuple[str, str]]:
    """
    Return (matched_keyword, product_text) for the first keyword found in a
    CDS feature's qualifier text, or None if no keyword matches.

    Parameters
    ----------
    feature : SeqFeature
        A Biopython CDS SeqFeature.
    keywords : tuple[str, ...]
        Keywords to test (case-insensitive substring match).

    Returns
    -------
    tuple[str, str] or None
        (matched_keyword, product_annotation_text) on the first hit.
    """
    text = get_qualifier_text(feature)
    for kw in keywords:
        if kw in text:
            product = feature.qualifiers.get("product", ["(no product qualifier)"])[0]
            return kw, product
    return None


def detect_holin_and_rbp(record: SeqRecord) -> dict:
    """
    Scan a SeqRecord's CDS features for holin and tail fiber/RBP.

    For each category, the FIRST matching CDS is recorded together with the
    keyword that matched and the product annotation text, so that every
    "Yes" call is auditable against the source GenBank record.

    Parameters
    ----------
    record : SeqRecord
        A Biopython SeqRecord parsed from a GenBank flat file.

    Returns
    -------
    dict
        Keys: Phage, Accession,
              Holin_Present, Holin_Evidence,
              Tail_Fiber_RBP_Present, RBP_Evidence,
              Annotation_Note.
        *_Evidence columns show "[matched_keyword] product_text" for each Yes,
        or "—" for a No. These columns make the detection verifiable and can
        be dropped when assembling the final Table 2.
    """
    holin_evidence: Optional[tuple[str, str]] = None
    rbp_evidence:   Optional[tuple[str, str]] = None

    for feature in record.features:
        if feature.type != "CDS":
            continue

        if holin_evidence is None:
            holin_evidence = _match_keyword(feature, HOLIN_KEYWORDS)

        if rbp_evidence is None:
            rbp_evidence = _match_keyword(feature, RBP_KEYWORDS)

        # Early exit once both are found
        if holin_evidence is not None and rbp_evidence is not None:
            break

    note = (
        "Detection by GenBank text annotation only. "
        "'No' may reflect incomplete annotation, not biological absence."
    )

    def fmt(ev: Optional[tuple[str, str]]) -> str:
        return f"[{ev[0]}] {ev[1]}" if ev else "—"

    return {
        "Phage":                  record.annotations.get("organism", record.name),
        "Accession":              record.id,
        "Holin_Present":          "Yes" if holin_evidence else "No",
        "Holin_Evidence":         fmt(holin_evidence),
        "Tail_Fiber_RBP_Present": "Yes" if rbp_evidence   else "No",
        "RBP_Evidence":           fmt(rbp_evidence),
        "Annotation_Note":        note,
    }


def parse_file(gb_path: Path) -> Optional[dict]:
    """Parse one GenBank file and return its holin/RBP detection result."""
    try:
        record = SeqIO.read(str(gb_path), "genbank")
        return detect_holin_and_rbp(record)
    except ValueError as exc:
        log.warning(f"  Skipped '{gb_path.name}': {exc}")
    except Exception as exc:
        log.warning(f"  Skipped '{gb_path.name}': unexpected error — {exc}")
    return None


# ===========================================================================
# Batch processing and I/O
# ===========================================================================

def run(input_dir: Path, output_path: Path) -> pd.DataFrame:
    """
    Process all GenBank files and write the holin/RBP annotation table.

    Parameters
    ----------
    input_dir : Path
        Directory containing GenBank flat files.
    output_path : Path
        Output CSV file path (UTF-8 encoded).

    Returns
    -------
    pd.DataFrame
        Annotation table sorted by Accession.
    """
    gb_files = sorted(
        f for f in input_dir.iterdir()
        if f.is_file() and f.suffix.lower() in VALID_EXTENSIONS
    )

    if not gb_files:
        log.error(f"No GenBank files found in '{input_dir}'.")
        sys.exit(1)

    log.info(f"Found {len(gb_files)} GenBank file(s)")

    rows = []
    for gb in gb_files:
        log.info(f"  Processing {gb.name}")
        row = parse_file(gb)
        if row:
            rows.append(row)

    if not rows:
        log.error("No records extracted.")
        sys.exit(1)

    df = pd.DataFrame(rows).sort_values("Accession").reset_index(drop=True)

    output_path.parent.mkdir(parents=True, exist_ok=True)
    df.to_csv(output_path, index=False, encoding="utf-8")

    log.info(f"Table 2 (partial) written: '{output_path}'  ({len(df)} records)")
    return df


# ===========================================================================
# CLI
# ===========================================================================

def build_parser() -> argparse.ArgumentParser:
    parser = argparse.ArgumentParser(
        prog="S2_holin_tailfiber_annotation.py",
        description=(
            "Keyword-based detection of Holin and Tail Fiber/RBP from "
            "Staphylococcus phage GenBank annotations. "
            "Generates Holin_Present and Tail_Fiber_RBP_Present columns for Table 2. "
            "Endolysin/domain data requires S4 + InterPro (separate workflow)."
        ),
        formatter_class=argparse.RawDescriptionHelpFormatter,
        epilog="""
MANUSCRIPT OUTPUT
-----------------
  TABLE 2 columns (partial):
    Holin Present | Tail Fiber / RBP Present

  Remaining Table 2 columns require:
    → Run S4_endolysin_extractor_for_interpro.py
    → Submit .faa files to InterPro (https://www.ebi.ac.uk/interpro/)

Examples (Windows Command Prompt)
----------------------------------
  python S2_holin_tailfiber_annotation.py -i GenBank -o results\\Table2_holin_rbp.csv
        """,
    )
    parser.add_argument(
        "--input_dir", "-i",
        type=Path, required=True, metavar="DIR",
        help="Directory containing GenBank files (.gb / .gbk / .gbff)",
    )
    parser.add_argument(
        "--output", "-o",
        type=Path, default=Path("Table2_holin_rbp.csv"), metavar="FILE",
        help="Output CSV file path. Default: Table2_holin_rbp.csv",
    )
    return parser


# ===========================================================================
# Entry point
# ===========================================================================

if __name__ == "__main__":
    args = build_parser().parse_args()
    df   = run(args.input_dir, args.output)

    sep = "=" * 68
    print(f"\n{sep}")
    print("TABLE 2 (PARTIAL) — HOLIN AND TAIL FIBER/RBP DETECTION")
    print(sep)
    print(df[["Accession", "Holin_Present", "Tail_Fiber_RBP_Present", "RBP_Evidence"]].to_string(index=False))
    print(sep)
    print(f"  Genomes analysed         : {len(df)}")
    print(f"  Holin detected (Yes)     : {(df['Holin_Present'] == 'Yes').sum()}")
    print(f"  Tail fiber/RBP detected  : {(df['Tail_Fiber_RBP_Present'] == 'Yes').sum()}")
    print(sep)
    print("\nREMINDER: For complete Table 2 (endolysin, domains),")
    print("run S4_endolysin_extractor_for_interpro.py then submit to InterPro.")
```
