## Supplementary material for "In silico characterization of lysis and host-recognition modules in *Staphylococcus aureus* bacteriophage genomes": https://github.com/HivanA98/In-silico-characterization-of-lysis-and-host-recognition-modules-in-S.-aureus-bacteriophage-genomes: S3_terl_extractor.py

```
#!/usr/bin/env python3
"""
S3_terl_extractor.py
====================
Terminase Large Subunit (TerL) Extractor for Phylogenetic Analysis

==============================================================================
MANUSCRIPT CONTRIBUTION
  Generates input for:  FIGURE 1 — Maximum Likelihood Phylogenetic Tree

  Complete Figure 1 pipeline:
    Step 1 — This script   → TerL_combined.faa  (multi-FASTA)
    Step 2 — MAFFT         → TerL_aligned.faa   (multiple sequence alignment)
    Step 3 — MEGA 12.1.2   → Figure 1           (Maximum Likelihood tree)
  See "Downstream Phylogenetic Workflow" below for exact commands and MEGA
  parameter settings.
==============================================================================

Associated manuscript:
    "Molecular Characterization of Lytic Bacteriophages Against Resistant
    Staphylococcus aureus Based on NCBI GenBank Sequences:
    A Bioinformatic Literature Review"

Description
-----------
Extracts terminase large subunit (TerL) amino acid sequences from a directory
of GenBank files and writes a single combined multi-FASTA file for direct
alignment and phylogenetic analysis.

TerL is the standard phylogenetic marker for tailed bacteriophage
classification: universally present, under strong purifying selection,
and provides clear inter-family resolution (Meier-Kolthoff & Goker, 2017).

Historical Background and Annotation Problem Resolved
------------------------------------------------------
During the original analysis, the extraction script was initially unable to
identify TerL in 9 of 22 genomes. Investigation revealed two distinct
situations:

  Case A — Kayvirus group (7 genomes: NC_047722, NC_047723, NC_047724,
  NC_047725, NC_047726, NC_047727, EU418428):
    The TerL protein in these genomes (605 aa) is annotated in the GenBank
    "product" qualifier with the 3-letter abbreviation "Ter" rather than the
    standard text "terminase large subunit". This non-standard annotation
    causes a failure in standard keyword searches.
    FIX: EXACT_NAMES = {"Ter", "ter"} — checks if the product qualifier
    EXACTLY EQUALS "Ter" or "ter" (case-sensitive). This catches all 7
    Kayvirus genomes and is the critical correction in this script.

  Case B — Staphylococcus phage Portland (MT926124) and
  Staphylococcus phage vB_SauP-436A1 (MN150710):
    These two genomes genuinely have no annotated TerL sequence. Detailed
    inspection reveals:
      Portland (MT926124):        "putative encapsidation protein" (415 aa)
      vB_SauP-436A1 (MN150710):  "DNA packaging protein" (415 aa)
    Both are micro-phages (~17–18 kb genome) that likely do not belong to
    Myovirus morphology and use a packaging mechanism not annotated under
    standard TerL nomenclature. Exclusion from phylogenetic analysis is
    scientifically justified.
    DECISION: These two genomes are CORRECTLY EXCLUDED from the output FASTA
    and from Figure 1.

Methods Statement for Manuscript
---------------------------------
The following text was incorporated into the Methods section:

  "Staphylococcus phage Portland (MT926124) and vB_SauP-436A1 (MN150710)
  were excluded from phylogenetic analysis due to the absence of annotated
  terminase large subunit sequences, consistent with their atypical small
  genome sizes (<20 kb) relative to the remaining dataset."

Detection Logic (Two Mechanisms)
----------------------------------
For each CDS feature in a GenBank record, two checks are applied:

  Mechanism 1 — Keyword match:
    Any string from TERL_KEYWORDS is found as a substring in the lowercased
    "product" qualifier. Catches standard annotations such as
    "terminase large subunit", "large terminase", "TerL protein", etc.

  Mechanism 2 — Exact product match:
    The product qualifier EXACTLY EQUALS "Ter" or "ter" (case-sensitive,
    checked against EXACT_NAMES). This is the fix for the Kayvirus group.

The FIRST CDS feature satisfying either mechanism AND containing a valid
"translation" qualifier is returned as the TerL for that genome.

Output FASTA Header Format
--------------------------
    >{accession}|{organism_no_spaces}|{product_annotation}
Example:
    >NC_047722.1|Staphylococcus_phage_Staph1N|Ter

This format is directly importable by MAFFT and MEGA 12.1.2.

Downstream Phylogenetic Workflow (Figure 1)
-------------------------------------------
Step 1 — Run this script:
    python S3_terl_extractor.py -i GenBank -o results\\TerL_combined.faa

Step 2 — Multiple Sequence Alignment with MAFFT (WEB SERVER, not local app):
    Open the MAFFT online server:
      https://mafft.cbrc.jp/alignment/server/

    a. Upload results\\TerL_combined.faa, OR paste the FASTA content into
       the input box.
    b. Under "Advanced settings", set the alignment strategy to:
         L-INS-i
       (Very slow; recommended for <200 sequences with one conserved
        domain and long gaps; 2 iterative cycles only.)
       This is the strategy used for the manuscript: TerL is a single
       conserved domain, and L-INS-i gives the most accurate alignment
       for a dataset of this size (20 sequences).
    c. Click "Submit".
    d. When alignment completes, open the "Fasta format" result and save
       it as results\\TerL_aligned.faa (use "Save As" in the browser, or
       copy the FASTA text into a new file).

Step 3 — Phylogenetic Tree in MEGA 12.1.2:
    Download MEGA 12.1.2: https://www.megasoftware.net/

    Open results\\TerL_aligned.faa in MEGA 12.1.2, then:
      Menu: Phylogeny > Construct/Test Maximum Likelihood Tree
        Substitution model    : LG+G+I
        Rates among sites     : Gamma distribution + Invariant sites (G+I)
        ML heuristic method   : Nearest-Neighbor-Interchange (NNI)
        Bootstrap replicates  : 1000
        Partial deletion      : 80%% site coverage cutoff
        Gaps/Missing data     : Partial Deletion
        Outgroup              : Staphylococcus phage EW (NC_007056.1)
        Condense tree at      : 50%% bootstrap (for polytomy display)

Reproducibility Note (tree topology vs bootstrap values)
----------------------------------------------------------
The TREE TOPOLOGY (branching pattern) is the reportable, stable result and
is reproducible given the same input sequences and the same MAFFT strategy
(L-INS-i). However, the BOOTSTRAP SUPPORT VALUES will vary by a few percent
between runs even on an identical alignment, because bootstrapping is a
random-resampling procedure and MEGA uses a different random seed each run.
A change such as 30 -> 26 or 50 -> 47 at a node reflects this normal
stochastic variation, NOT a change in the underlying phylogeny. To minimise
run-to-run differences, always use the same MAFFT strategy (L-INS-i) and the
same 1000-replicate / 80%%-partial-deletion settings; report the topology and
the approximate support, not exact bootstrap integers.

Reference
---------
Cock PJA, Antao T, Chang JT, Chapman BA, Cox CJ, Dalke A, Friedberg I,
Hamelryck T, Kauff F, Wilczynski B, de Hoon MJL (2009). Biopython: freely
available Python tools for computational molecular biology and bioinformatics.
Bioinformatics, 25(11):1422-1423. doi:10.1093/bioinformatics/btp163

Meier-Kolthoff JP, Goker M (2017). VICTOR: genome-based phylogeny and
classification of prokaryotic viruses. Bioinformatics, 33(21):3396-3404.
doi:10.1093/bioinformatics/btx440

Tested Environment (Windows)
-----------------------------
    OS         : Windows 10 / 11
    Python     : 3.12.10
    biopython  : 1.87
    MAFFT      : WEB SERVER (https://mafft.cbrc.jp/alignment/server/),
                 strategy L-INS-i — no local MAFFT installation required
    MEGA       : 12.1.2 (https://www.megasoftware.net)

Installation (Command Prompt / PowerShell)
------------------------------------------
    pip install biopython==1.87

Usage (Windows Command Prompt)
-------------------------------
    python S3_terl_extractor.py -i GenBank -o results\\TerL_combined.faa
"""

import argparse
import logging
import sys
from dataclasses import dataclass
from pathlib import Path
from typing import Optional

from Bio import SeqIO
from Bio.SeqRecord import SeqRecord

# ---------------------------------------------------------------------------
# Logging
# ---------------------------------------------------------------------------
logging.basicConfig(
    level=logging.INFO,
    format="%(asctime)s  [%(levelname)-8s]  %(message)s",
    datefmt="%Y-%m-%d %H:%M:%S",
)
log = logging.getLogger(__name__)

# ---------------------------------------------------------------------------
# Constants
# ---------------------------------------------------------------------------
VALID_EXTENSIONS: frozenset[str] = frozenset({".gb", ".gbk", ".gbff"})

# ---- Mechanism 1: Keyword match on "product" qualifier ----
# Matched case-insensitively as substrings.
TERL_KEYWORDS: tuple[str, ...] = (
    "terminase large subunit",
    "large terminase",
    "large subunit terminase",
    "terminase, large subunit",
    "dna packaging terminase large subunit",
    "dna terminase large subunit",
    "large terminase subunit",
    "terl",                  # catches "TerL", "TerL protein", etc.
)

# ---- Mechanism 2: Exact match on "product" qualifier ----
# Checks if the product annotation is EXACTLY one of these strings
# (case-sensitive). Required to catch Kayvirus group annotation "Ter".
EXACT_NAMES: frozenset[str] = frozenset({
    "Ter",    # Kayvirus group (NC_047722–727 and EU418428): TerL is 605 aa,
    "ter",    # annotated as product="Ter" — non-standard 3-letter abbreviation.
})


# ===========================================================================
# Data container
# ===========================================================================

@dataclass
class TerLRecord:
    """Container for an extracted TerL sequence and detection metadata."""
    accession:         str
    organism:          str
    product:           str
    sequence:          str
    detection_method:  str    # "keyword_match" or "exact_product_match"

    @property
    def length(self) -> int:
        return len(self.sequence)

    @property
    def fasta_header(self) -> str:
        """
        FASTA header: {accession}|{organism_no_spaces}|{product}

        The detection_method is NOT included in the header to keep the
        FASTA file compatible with MAFFT and MEGA sequence labeling.
        """
        org_safe  = self.organism.replace(" ", "_")
        prod_safe = self.product.replace("|", "/")
        return f"{self.accession}|{org_safe}|{prod_safe}"


# ===========================================================================
# Detection logic
# ===========================================================================

def find_terl_in_record(record: SeqRecord) -> Optional[TerLRecord]:
    """
    Search all CDS features in a SeqRecord for the TerL sequence.

    Applies Mechanism 1 (keyword match) then Mechanism 2 (exact product match)
    to each CDS feature. Returns the FIRST feature that:
      (a) satisfies either mechanism, AND
      (b) contains a non-empty "translation" qualifier.

    Parameters
    ----------
    record : SeqRecord
        A Biopython SeqRecord from a GenBank flat file.

    Returns
    -------
    TerLRecord or None
        Extracted TerL record on success; None if no TerL CDS is found
        or all matching CDS lack a translation qualifier.

    Notes
    -----
    Bacteriophage genomes encode one terminase large subunit. Multiple matches
    would indicate annotation redundancy rather than biology; only the first
    valid match is returned.
    """
    organism = record.annotations.get("organism", record.name)

    for feature in record.features:
        if feature.type != "CDS":
            continue

        product_text = " ".join(feature.qualifiers.get("product", [])).lower()
        product_raw  = feature.qualifiers.get("product", ["terminase large subunit"])[0]

        # Mechanism 1: keyword substring match
        if any(kw in product_text for kw in TERL_KEYWORDS):
            method = "keyword_match"
        # Mechanism 2: exact product name match (catches Kayvirus "Ter")
        elif product_raw in EXACT_NAMES:
            method = "exact_product_match"
        else:
            continue

        # Verify translation qualifier exists
        translation = feature.qualifiers.get("translation", [])
        if not translation or not translation[0]:
            log.warning(
                f"  {record.id}: TerL match ('{product_raw}', {method}) "
                f"has no 'translation' qualifier — skipped"
            )
            continue

        return TerLRecord(
            accession=record.id,
            organism=organism,
            product=product_raw,
            sequence=translation[0],
            detection_method=method,
        )

    return None


def parse_file(gb_path: Path) -> Optional[TerLRecord]:
    """Parse one GenBank file and extract its TerL sequence."""
    try:
        record = SeqIO.read(str(gb_path), "genbank")
        return find_terl_in_record(record)
    except ValueError as exc:
        log.warning(f"  Skipped '{gb_path.name}': {exc}")
    except Exception as exc:
        log.warning(f"  Skipped '{gb_path.name}': unexpected error — {exc}")
    return None


# ===========================================================================
# Batch processing and I/O
# ===========================================================================

def run(input_dir: Path, output_path: Path) -> list[TerLRecord]:
    """
    Process all GenBank files and write combined multi-FASTA for MAFFT/MEGA.

    Genomes without a detectable TerL (Portland MT926124 and
    vB_SauP-436A1 MN150710) are excluded from the FASTA output and
    reported in the terminal log, consistent with their exclusion from
    Figure 1 in the manuscript.

    Parameters
    ----------
    input_dir : Path
        Directory containing GenBank flat files.
    output_path : Path
        Output multi-FASTA file (.faa).

    Returns
    -------
    list[TerLRecord]
        Extracted TerL records, sorted by accession.
    """
    gb_files = sorted(
        f for f in input_dir.iterdir()
        if f.is_file() and f.suffix.lower() in VALID_EXTENSIONS
    )

    if not gb_files:
        log.error(f"No GenBank files found in '{input_dir}'.")
        sys.exit(1)

    log.info(f"Found {len(gb_files)} GenBank file(s) in '{input_dir}'")

    found:     list[TerLRecord] = []
    not_found: list[str]        = []

    for gb in gb_files:
        log.info(f"  Processing {gb.name}")
        terl = parse_file(gb)
        if terl:
            found.append(terl)
            log.info(
                f"    ✓  {terl.accession:<16}  {terl.length:>4} aa  "
                f"[{terl.detection_method}]  product: '{terl.product}'"
            )
        else:
            not_found.append(gb.stem)   # accession stem
            log.info(f"    ✗  {gb.stem}  — no TerL detected (correctly excluded)")

    if not found:
        log.error("No TerL sequences extracted. Check GenBank annotations.")
        sys.exit(1)

    # Write combined multi-FASTA sorted by accession
    output_path.parent.mkdir(parents=True, exist_ok=True)
    with open(output_path, "w", encoding="utf-8") as fh:
        for t in sorted(found, key=lambda x: x.accession):
            fh.write(f">{t.fasta_header}\n{t.sequence}\n")

    log.info(f"\nOutput multi-FASTA written: '{output_path}'")
    log.info(f"  Sequences included : {len(found)}")
    log.info(f"  Genomes excluded   : {len(not_found)}")

    if not_found:
        log.info("  Excluded genomes (no annotated TerL):")
        for acc in not_found:
            log.info(f"    • {acc}")
        log.info(
            "  Methods statement: 'Staphylococcus phage Portland (MT926124) "
            "and vB_SauP-436A1 (MN150710) were excluded from phylogenetic "
            "analysis due to the absence of annotated terminase large subunit "
            "sequences, consistent with their atypical small genome sizes "
            "(<20 kb) relative to the remaining dataset.'"
        )

    return found


# ===========================================================================
# CLI
# ===========================================================================

def build_parser() -> argparse.ArgumentParser:
    parser = argparse.ArgumentParser(
        prog="S3_terl_extractor.py",
        description=(
            "Extract TerL sequences from Staphylococcus phage GenBank files. "
            "Applies two detection mechanisms: (1) keyword match and "
            "(2) exact product match for 'Ter'/'ter' (Kayvirus group fix). "
            "Outputs combined multi-FASTA for MAFFT + MEGA 12.1.2 (Figure 1)."
        ),
        formatter_class=argparse.RawDescriptionHelpFormatter,
        epilog="""
MANUSCRIPT OUTPUT
-----------------
  FIGURE 1 — Maximum Likelihood Phylogenetic Tree

  Pipeline for Figure 1:
    Step 1: python S3_terl_extractor.py -i GenBank -o results\\TerL_combined.faa
    Step 2: Align on MAFFT web server (https://mafft.cbrc.jp/alignment/server/)
            Upload TerL_combined.faa, set strategy = L-INS-i, Submit,
            save result as TerL_aligned.faa
    Step 3: Open results\\TerL_aligned.faa in MEGA 12.1.2
            Phylogeny > ML > LG+G+I > 1000 bootstrap > 80%% partial deletion
            Outgroup: Staphylococcus phage EW (NC_007056.1)

Detection mechanisms:
  Mechanism 1: keyword match in product qualifier (standard annotations)
  Mechanism 2: exact match for product="Ter"/"ter" (Kayvirus group fix)

Genomes excluded from FASTA (no annotated TerL; consistent with Figure 1):
  • Portland (MT926124)     — 17,471 bp, no TerL annotation
  • vB_SauP-436A1 (MN150710) — 18,028 bp, no TerL annotation

Examples (Windows Command Prompt)
----------------------------------
  python S3_terl_extractor.py -i GenBank -o results\\TerL_combined.faa
        """,
    )
    parser.add_argument(
        "--input_dir", "-i",
        type=Path, required=True, metavar="DIR",
        help="Directory containing GenBank files (.gb / .gbk / .gbff)",
    )
    parser.add_argument(
        "--output", "-o",
        type=Path, default=Path("TerL_combined.faa"), metavar="FILE",
        help="Output multi-FASTA file. Default: TerL_combined.faa",
    )
    return parser


# ===========================================================================
# Entry point
# ===========================================================================

if __name__ == "__main__":
    args  = build_parser().parse_args()
    found = run(args.input_dir, args.output)

    sep = "=" * 80
    kw_count    = sum(1 for t in found if t.detection_method == "keyword_match")
    exact_count = sum(1 for t in found if t.detection_method == "exact_product_match")

    print(f"\n{sep}")
    print("FIGURE 1 INPUT — TerL EXTRACTION SUMMARY")
    print(sep)
    print(f"{'Accession':<20} {'Organism':<36} {'Length':>6}  {'Method':<22}  {'Product'}")
    print("-" * 80)
    for t in sorted(found, key=lambda x: x.accession):
        print(f"{t.accession:<20} {t.organism[:35]:<36} {t.length:>6}  "
              f"{t.detection_method:<22}  {t.product}")
    print(sep)
    print(f"  Total sequences extracted : {len(found)}")
    print(f"  Via keyword_match         : {kw_count}  (standard annotations)")
    print(f"  Via exact_product_match   : {exact_count}  (Kayvirus 'Ter' fix)")
    print(f"\n  Output FASTA   : {args.output}")
    print(f"\nNEXT STEP:")
    print(f"  1. Align on MAFFT web: https://mafft.cbrc.jp/alignment/server/")
    print(f"     Upload {args.output}, set strategy = L-INS-i, Submit,")
    print(f"     save result as TerL_aligned.faa")
    print(f"  2. Open TerL_aligned.faa in MEGA 12.1.2 -> ML tree")
    print(f"     (LG+G+I, 1000 bootstrap, outgroup EW NC_007056.1)")
```
