## Supplementary material for "In silico characterization of lysis and host-recognition modules in *Staphylococcus aureus* bacteriophage genomes": https://github.com/HivanA98/In-silico-characterization-of-lysis-and-host-recognition-modules-in-S.-aureus-bacteriophage-genomes: S4_endolysin_extractor_for_interpro.py

```
#!/usr/bin/env python3
"""
S4_endolysin_extractor_for_interpro.py
=======================================
Endolysin Classifier & Extractor — domain-aware, with tBLASTn fallback

==============================================================================
MANUSCRIPT CONTRIBUTION
  Completes the lysis-enzyme columns of:
    TABLE 2 — Endolysin gene product | Endolysin length (aa) |
              Catalytic domains | Wall-binding domain (CBD)

  WHY THIS VERSION REPLACES THE NAME-ONLY EXTRACTOR
  ──────────────────────────────────────────────────
  Selecting an endolysin by the FIRST product-name keyword match is unsafe.
  Manual InterPro + tBLASTn validation of this dataset exposed three failure
  modes that a name-only, first-match extractor gets wrong:

    Maine (MN045228)  : first match was the non-lytic "N-glycosidase
                        YbiA-like" protein; the real endolysin is a free
                        LysK recovered only by tBLASTn (99% identity to the
                        Sb1_8383 LysK reference).
    JD007 (NC_019726) : first match was a 295-aa virion-associated NlpC/P60
                        protein; the real endolysin is a free 495-aa LysK,
                        again recovered by tBLASTn (99% identity), missed by
                        the keyword scan.
    Twort (NC_007021) : first match was a 1269-aa phage tail lysozyme
                        (Phage_lysozyme2) — a virion-associated peptidoglycan
                        hydrolase (VAPH), NOT a free endolysin; tBLASTn finds
                        only a divergent (~45%) LysK homolog.

  This version therefore (1) collects ALL lysis-keyword CDS and ranks them,
  (2) infers a domain class from the annotation and flags non-lytic hits,
  (3) separates free endolysins from virion-associated (VAPH) enzymes,
  (4) runs an automatic tBLASTn fallback against a LysK reference when no
  valid free endolysin is found by keyword, (5) flags the known intron-split
  and HNH-disrupted ORFs, and (6) writes an auditable CSV plus a combined
  FASTA for InterPro.

  HONEST SCOPE OF "DOMAIN VALIDATION"
  ────────────────────────────────────
  A GenBank flat file carries product NAMES, not Pfam/InterPro domain
  assignments. True domain assignment therefore still comes from InterPro
  (https://www.ebi.ac.uk/interpro/search/sequence/) on the FASTA this script
  produces — that is the confirmatory step. Offline, this script INFERS a
  domain class from the product/gene/note text and from protein length, and
  FLAGS each candidate accordingly. The CSV has a blank "InterPro_Domain"
  column to be filled from the InterPro result. tBLASTn is wired as a real
  subprocess call to local BLAST+ (tblastn) when it is on PATH; otherwise the
  exact command is printed and the validated identity (below) is reported.

  AUTHORITATIVE VALIDATED RESULTS (KNOWN_CASES)
  ───────────────────────────────────────────────
  The classifications that required manual InterPro + tBLASTn work are encoded
  in KNOWN_CASES, taken directly from the validated Table 2 of the manuscript
  (Hasugian, Journal 2). For these accessions the script reports the validated
  status and tBLASTn identity rather than relying on fragile name heuristics.
  Any genome NOT in KNOWN_CASES is handled by the general heuristic logic, so
  the script also works on new inputs.
==============================================================================

Associated manuscript:
    "Molecular Characterization of Lytic Bacteriophages Against Resistant
    Staphylococcus aureus Based on NCBI GenBank Sequences:
    A Bioinformatic Literature Review"

Outputs
-------
  1. Combined FASTA (--output): one representative protein per genome where a
     sequence is available from the GenBank annotation (free endolysin,
     intron-split moiety, VAPH, or divergent endolysin), each header tagged
     with its classification. Sized for a single InterPro submission (<100).
  2. Audit CSV (--csv): one row per candidate CDS, with the matched product,
     inferred domain, classification, whether it was selected, tBLASTn
     identity/coordinates when used, and a free-text Evidence field.

Reference
---------
Cock PJA, et al. (2009). Biopython. Bioinformatics, 25(11):1422-1423.
Kornienko M, et al. (2020). Sci Rep / Viruses (Transcriptional Landscape, TU16).
Kornienko M, et al. (2023). Sb-1 lineage comparative genomics.

Dependencies (exact versions; no other dependencies required)
--------------------------------------------------------------
  pip:    biopython 1.87        GenBank parsing (Cock et al., 2009)
  stdlib: argparse, csv, logging, shutil, subprocess, sys, tempfile,
          dataclasses, pathlib, typing
  Output is plain-text FASTA and CSV (no Excel/openpyxl).
  Optional external tool: NCBI BLAST+ (tblastn) for the live tBLASTn fallback.

Tested Environment (Windows)
-----------------------------
    OS        : Windows 10 / 11
    Python    : 3.12.10
    biopython : 1.87
    BLAST+    : optional (tblastn on PATH) — else the command is printed

Usage (Windows Command Prompt)
-------------------------------
    python S4_endolysin_extractor_for_interpro.py -i GenBank ^
        -o results\\endolysin_candidates.faa --csv results\\endolysin_audit.csv

    # enable the live tBLASTn fallback (requires BLAST+):
    python S4_endolysin_extractor_for_interpro.py -i GenBank ^
        -o results\\endolysin_candidates.faa --csv results\\endolysin_audit.csv ^
        --run-tblastn
"""

import argparse
import csv
import logging
import shutil
import subprocess
import sys
import tempfile
from dataclasses import dataclass, field
from pathlib import Path
from typing import Optional

from Bio import SeqIO
from Bio.SeqRecord import SeqRecord

# ---------------------------------------------------------------------------
# Logging
# ---------------------------------------------------------------------------
logging.basicConfig(
    level=logging.INFO,
    format="%(asctime)s  [%(levelname)-8s]  %(message)s",
    datefmt="%Y-%m-%d %H:%M:%S",
)
log = logging.getLogger(__name__)

# ---------------------------------------------------------------------------
# Constants
# ---------------------------------------------------------------------------
VALID_EXTENSIONS: frozenset[str] = frozenset({".gb", ".gbk", ".gbff"})
INTERPRO_MAX_SEQUENCES: int = 100

# (1) Broad collection keywords — match ANY potential lysis-module CDS so that
# non-lytic and VAPH proteins are also collected and then correctly classified.
LYSIS_KEYWORDS: tuple[str, ...] = (
    "lysin", "endolysin", "lysk", "amidase", "chap", "peptidase",
    "hydrolase", "muramidase", "glucosaminidase", "glycosidase",
    "lysozyme", "peptidoglycan", "nlpc", "p60",
)

# (2) Non-lytic markers — these are NOT peptidoglycan hydrolases. A candidate
# whose annotation carries one of these is flagged non-lytic and rejected as an
# endolysin. NOTE: plain "glycosidase" is NOT here, because a phosphodiester
# NAGPA glycosidase can be a genuine (divergent) endolysin (e.g., phage EW).
# Only the YbiA / NADAR family N-glycosidase is non-lytic.
NONLYTIC_MARKERS: tuple[str, ...] = (
    "ybia", "nadar",
)

# (3) VAPH (virion-associated peptidoglycan hydrolase) markers — tail/baseplate
# anchored structural hydrolases. A candidate carrying any of these is a VAPH,
# not a free endolysin.
VAPH_MARKERS: tuple[str, ...] = (
    "tail", "baseplate", "structural", "virion", "lid_weld",
    "phage_lysozyme2", "tail-anchored", "tail associated", "tail-associated",
)
# Backup length cut: free Staph endolysins in this dataset are <=576 aa;
# the only >700 aa lysis CDS is the 1269-aa Twort tail lysozyme (a VAPH).
VAPH_LENGTH_THRESHOLD_AA: int = 700

# Recognised peptidoglycan-hydrolase (lytic) domain terms used by the inference.
LYTIC_DOMAIN_TERMS: tuple[str, ...] = (
    "chap", "amidase", "n-acetylmuramoyl", "peptidase_m15", "m15",
    "glucosaminidase", "muramidase", "nlpc", "p60", "phage_lysozyme",
    "lysozyme", "lysk", "endolysin", "peptidoglycan hydrolase",
    "glycosidase",   # divergent endolysins (NAGPA/phosphodiester) — see EW
)

# (5) Known special cases — runtime flags.
INTRON_SPLIT_ACCESSIONS:   frozenset[str] = frozenset({"MN047438", "MF398190"})
HNH_FRAGMENTED_ACCESSIONS: frozenset[str] = frozenset({"MN336262", "MN336263"})

# Default tBLASTn reference: the Sb1_8383 LysK (accession MN336261), the query
# used in the manuscript's manual validation. Auto-extracted from the dataset
# if present; overridable with --reference.
DEFAULT_REFERENCE_ACCESSION: str = "MN336261"

# Authoritative validated results (from the manuscript's InterPro + tBLASTn
# Table 2). status, human-readable note, and tBLASTn % identity (or None).
# These OVERRIDE the heuristic for the listed accessions only.
KNOWN_CASES: dict[str, dict] = {
    "MN047438": dict(status="intron-split", identity=None,
        note="lysK.1 N-terminal amidase moiety (~209 aa); full LysK ~495 aa "
             "reconstructed across a self-splicing intron (Kornienko 2020, "
             "Viruses, TU16). Catalytic Amidase here; CHAP on C-terminal moiety; SH3b CBD."),
    "MF398190": dict(status="intron-split", identity=None,
        note="lysK.1 N-terminal amidase moiety (~209 aa); full LysK ~495 aa "
             "across intron (same architecture as MN047438)."),
    "MN045228": dict(status="tblastn-recovered", identity=99.0,
        note="free LysK missed by keyword scan; recovered by tBLASTn vs "
             "Sb1_8383 at 99% identity (full ~495 aa, CHAP+Amidase+SH3b)."),
    "NC_019726": dict(status="tblastn-recovered", identity=99.0,
        note="free 495-aa LysK recovered by tBLASTn vs Sb1_8383 at 99%; "
             "keyword scan returned a 295-aa virion-associated NlpC/P60 (VAPH)."),
    "MN336262": dict(status="hnh-disrupted", identity=None,
        note="no intact endolysin ORF (HNH endonuclease insertion); 141-aa "
             "CHAP-bearing fragment recovered by tBLASTn (c29476-29051); "
             "LysM CBD annotated separately (Kornienko 2023)."),
    "MN336263": dict(status="hnh-disrupted", identity=None,
        note="no intact endolysin ORF (HNH insertion); 141-aa CHAP fragment "
             "recovered by tBLASTn; LysM CBD annotated separately."),
    "NC_007021": dict(status="vaph", identity=45.0,
        note="1269-aa phage tail lysozyme (Phage_lysozyme2), tail-anchored — "
             "a virion-associated peptidoglycan hydrolase, NOT a free "
             "endolysin; only ~45% divergent LysK homology by tBLASTn."),
    "NC_007056": dict(status="divergent-endolysin", identity=None,
        note="576-aa NAGPA phosphodiester glycosidase, no SH3b CBD — a "
             "divergent free endolysin (outgroup lineage), not a non-lytic protein."),
}


# ---------------------------------------------------------------------------
# Data container
# ---------------------------------------------------------------------------

@dataclass
class Candidate:
    """One lysis-keyword CDS candidate from a genome."""
    protein_id: str
    accession:  str
    organism:   str
    product:    str
    sequence:   str
    inferred_domain: str = "unknown"
    classification:  str = "uncertain"   # free-endolysin|VAPH|non-lytic|intron-split|hnh-disrupted|tblastn-recovered|divergent-endolysin|uncertain
    selected:        bool = False
    tblastn_identity: Optional[float] = None
    tblastn_note:    str = ""
    runtime_flag:    str = ""
    evidence:        str = ""

    @property
    def length(self) -> int:
        return len(self.sequence)

    @property
    def fasta_header(self) -> str:
        org = self.organism.replace(" ", "_")
        prod = self.product.replace("|", "/")
        return f"{self.protein_id}|{self.accession}|{org}|status={self.classification}|{prod}"


# ===========================================================================
# Annotation parsing & inference
# ===========================================================================

def get_qualifier_text(feature) -> str:
    """Concatenate product/gene/note/function qualifiers, lowercased."""
    parts = []
    for key in ("product", "gene", "note", "function"):
        parts.extend(feature.qualifiers.get(key, []))
    return " ".join(parts).lower()


def infer_domain(text: str) -> str:
    """
    Infer a domain-class label from annotation text (lowercased).

    This is a NAME-BASED inference, confirmed later by InterPro. Order matters:
    the most specific / decisive markers are tested first.
    """
    if any(m in text for m in NONLYTIC_MARKERS):
        return "NADAR/YbiA (non-lytic)"
    if "phage_lysozyme2" in text or ("tail" in text and "lysozyme" in text):
        return "Phage_lysozyme2 (tail lysozyme)"
    has_chap = "chap" in text
    has_ami  = "amidase" in text or "n-acetylmuramoyl" in text
    if has_chap and has_ami:
        return "CHAP + Amidase (LysK-type)"
    if has_chap:
        return "CHAP"
    if has_ami:
        return "Amidase"
    if "m15" in text:
        return "Peptidase_M15"
    if "glucosaminidase" in text:
        return "Glucosaminidase"
    if "nlpc" in text or "p60" in text:
        return "NlpC/P60"
    if "muramidase" in text or "lysozyme" in text:
        return "Muramidase/Phage_lysozyme"
    if "glycosidase" in text:
        return "Glycosidase (phosphodiester/NAGPA)"
    if "lysk" in text or "endolysin" in text or "lysin" in text:
        return "LysK-type (unspecified)"
    if "peptidase" in text:
        return "Peptidase (unspecified)"
    if "hydrolase" in text or "peptidoglycan" in text:
        return "Peptidoglycan hydrolase (unspecified)"
    return "unknown"


def classify(accession: str, text: str, length: int, inferred: str) -> tuple[str, str]:
    """
    Classify a candidate by its OWN inferred nature, and assign any runtime flag.

    Returns (classification, runtime_flag). This is per-candidate and purely
    heuristic, so the audit CSV reflects each protein's true nature (e.g., a
    YbiA decoy is 'non-lytic', an NlpC/P60 virion protein is 'VAPH'). The
    genome-level validated status from KNOWN_CASES is applied later, only to
    the single representative chosen for that genome (see select_for_genome).
    """
    acc = accession.split(".")[0]
    runtime = ("intron-split" if acc in INTRON_SPLIT_ACCESSIONS else
               "hnh-disrupted" if acc in HNH_FRAGMENTED_ACCESSIONS else "")

    if any(m in inferred.lower() for m in ("non-lytic", "nadar", "ybia")):
        return "non-lytic", runtime
    is_vaph = any(m in text for m in VAPH_MARKERS) or length > VAPH_LENGTH_THRESHOLD_AA
    if is_vaph:
        return "VAPH", runtime
    is_lytic = any(t in inferred.lower() for t in
                   ("chap", "amidase", "m15", "glucosaminidase", "muramidase",
                    "nlpc", "p60", "lysozyme", "lysk", "glycosidase",
                    "peptidoglycan hydrolase"))
    if is_lytic:
        return "free-endolysin", runtime
    return "uncertain", runtime


def collect_candidates(record: SeqRecord) -> list[Candidate]:
    """Collect and classify ALL lysis-keyword CDS in a genome (requirement 1+2+3)."""
    organism  = record.annotations.get("organism", record.name)
    accession = record.id
    out: list[Candidate] = []

    for idx, feature in enumerate(record.features):
        if feature.type != "CDS":
            continue
        text = get_qualifier_text(feature)
        if not any(kw in text for kw in LYSIS_KEYWORDS):
            continue
        translation = feature.qualifiers.get("translation", [])
        if not translation or not translation[0]:
            continue

        product = feature.qualifiers.get("product", ["hypothetical protein"])[0]
        pid     = feature.qualifiers.get("protein_id", [f"CDS_{idx:04d}"])[0]
        seq     = translation[0]
        inferred = infer_domain(text)
        cls, flag = classify(accession, text, len(seq), inferred)

        out.append(Candidate(
            protein_id=pid, accession=accession, organism=organism,
            product=product, sequence=seq, inferred_domain=inferred,
            classification=cls, runtime_flag=flag,
        ))
    return out


def rank_score(c: Candidate) -> int:
    """Higher = stronger free-endolysin. Used to pick one primary per genome."""
    text = (c.product + " " + c.inferred_domain).lower()
    score = 0
    if c.classification == "free-endolysin":
        score += 1000
    if "endolysin" in text or "lysk" in text or "lysin" in text:
        score += 100
    if "chap" in text:
        score += 30
    if "amidase" in text:
        score += 30
    if 440 <= c.length <= 520:        # canonical full LysK
        score += 50
    elif 200 <= c.length <= 600:      # plausible endolysin
        score += 20
    if c.classification == "VAPH":
        score -= 500
    if c.classification == "non-lytic":
        score -= 1000
    return score


# ===========================================================================
# tBLASTn fallback (requirement 4)
# ===========================================================================

def parse_tblastn_outfmt6(text: str) -> Optional[dict]:
    """
    Parse BLAST -outfmt 6 (qseqid sseqid pident length sstart send evalue bitscore).
    Returns the best hit (highest bitscore) as a dict, or None if empty.
    """
    best = None
    for line in text.strip().splitlines():
        cols = line.split("\t")
        if len(cols) < 8:
            continue
        try:
            hit = dict(
                pident=float(cols[2]), length=int(cols[3]),
                sstart=int(cols[4]), send=int(cols[5]),
                evalue=float(cols[6]), bitscore=float(cols[7]),
            )
        except ValueError:
            continue
        if best is None or hit["bitscore"] > best["bitscore"]:
            best = hit
    return best


def find_tblastn() -> Optional[str]:
    """Return the tblastn executable path if BLAST+ is installed, else None."""
    return shutil.which("tblastn")


def build_reference(input_dir: Path, override: Optional[Path]) -> Optional[Path]:
    """
    Resolve the LysK reference FASTA for tBLASTn.

    Uses --reference if given; otherwise auto-extracts the best free endolysin
    from the Sb1_8383 record (DEFAULT_REFERENCE_ACCESSION, MN336261) if present.
    Returns a path to a FASTA, or None if no reference can be built.
    """
    if override is not None and override.is_file():
        log.info(f"  tBLASTn reference: {override} (user-provided)")
        return override

    for gb in input_dir.iterdir():
        if gb.suffix.lower() not in VALID_EXTENSIONS:
            continue
        if DEFAULT_REFERENCE_ACCESSION not in gb.stem:
            continue
        try:
            record = SeqIO.read(str(gb), "genbank")
        except Exception:
            continue
        cands = [c for c in collect_candidates(record)
                 if c.classification in ("free-endolysin", "tblastn-recovered")]
        if not cands:
            cands = collect_candidates(record)
        if not cands:
            return None
        ref_cand = max(cands, key=rank_score)
        tmp = Path(tempfile.gettempdir()) / "S4_lysK_reference.faa"
        tmp.write_text(f">Sb1_8383_LysK_reference|{ref_cand.accession}\n{ref_cand.sequence}\n",
                       encoding="utf-8")
        log.info(f"  tBLASTn reference: auto-extracted Sb1_8383 LysK "
                 f"({ref_cand.length} aa) from {gb.name}")
        return tmp

    log.info("  tBLASTn reference: not available "
             "(no MN336261 in input and no --reference given).")
    return None


def run_tblastn(record: SeqRecord, reference: Path, identity_threshold: float
                ) -> tuple[Optional[float], str, str]:
    """
    Run tBLASTn (LysK reference vs this genome) and interpret the best hit.

    Returns (identity_pct, note, status_suffix). If BLAST+ is unavailable, the
    ready-to-run command is returned in the note and identity is None.
    """
    exe = find_tblastn()
    if exe is None:
        cmd = (f'tblastn -query "{reference}" -subject <genome.fasta> '
               f'-outfmt "6 qseqid sseqid pident length sstart send evalue bitscore"')
        return None, f"BLAST+ not on PATH — run manually: {cmd}", "tblastn-required"

    with tempfile.NamedTemporaryFile("w", suffix=".fna", delete=False,
                                     encoding="utf-8") as fh:
        fh.write(f">{record.id}\n{str(record.seq)}\n")
        subject = fh.name

    try:
        proc = subprocess.run(
            [exe, "-query", str(reference), "-subject", subject,
             "-outfmt", "6 qseqid sseqid pident length sstart send evalue bitscore",
             "-max_target_seqs", "5"],
            capture_output=True, text=True, timeout=300,
        )
    except (subprocess.SubprocessError, OSError) as exc:
        return None, f"tblastn execution failed: {exc}", "tblastn-error"
    finally:
        Path(subject).unlink(missing_ok=True)

    hit = parse_tblastn_outfmt6(proc.stdout)
    if hit is None:
        return None, "tblastn returned no hit", "tblastn-no-hit"

    coords = f"c{hit['sstart']}-{hit['send']}" if hit["sstart"] > hit["send"] \
        else f"{hit['sstart']}-{hit['send']}"
    if hit["pident"] >= identity_threshold:
        return hit["pident"], (f"tBLASTn {hit['pident']:.1f}% over {hit['length']} aa "
                               f"at {coords} — free endolysin recovered"), "tblastn-recovered"
    return hit["pident"], (f"tBLASTn only {hit['pident']:.1f}% over {hit['length']} aa "
                           f"at {coords} — divergent/uncertain"), "divergent-uncertain"


# ===========================================================================
# Per-genome selection
# ===========================================================================

def select_for_genome(cands: list[Candidate], accession: str,
                      reference: Optional[Path], run_tblastn_flag: bool,
                      identity_threshold: float) -> tuple[Optional[Candidate], list[str]]:
    """
    Choose the representative endolysin for one genome and resolve special cases.

    Returns (selected_candidate_or_None, list_of_log_messages).
    """
    msgs: list[str] = []
    acc = accession.split(".")[0]

    # --- Authoritative validated cases ---
    if acc in KNOWN_CASES:
        kc = KNOWN_CASES[acc]
        status = kc["status"]
        # Pick the best available annotated sequence to represent this genome.
        if cands:
            if status == "vaph":
                pick = max(cands, key=lambda c: c.length)            # the tail lysozyme
            elif status == "intron-split":
                pick = max(cands, key=rank_score)                    # lysK.1 moiety
            elif status == "divergent-endolysin":
                pick = max(cands, key=rank_score)                    # the glycosidase
            else:  # tblastn-recovered / hnh-disrupted: annotation has only decoys/fragments
                pick = None
        else:
            pick = None

        if pick is not None:
            pick.classification = status
            pick.tblastn_identity = kc["identity"]
            pick.tblastn_note = kc["note"]
            pick.evidence = (f"matched '{pick.inferred_domain}'; validated status "
                             f"'{status}' (manuscript Table 2)")
            pick.selected = True
        # Report any tBLASTn identity from the validated record (and optionally re-run).
        ident_txt = f"{kc['identity']:.0f}%" if kc["identity"] is not None else "n/a"
        msgs.append(f"[{status}] {accession}: {kc['note']} (tBLASTn identity: {ident_txt})")
        if run_tblastn_flag and reference is not None and status in (
                "tblastn-recovered", "hnh-disrupted", "vaph"):
            ident, note, _ = run_tblastn_helper(acc, reference, identity_threshold)
            if ident is not None:
                msgs.append(f"    live tBLASTn re-check {accession}: {note}")
        return pick, msgs

    # --- General heuristic ---
    free = [c for c in cands if c.classification == "free-endolysin"]
    if free:
        pick = max(free, key=rank_score)
        pick.selected = True
        pick.evidence = (f"matched '{pick.product[:30]}'; inferred "
                         f"{pick.inferred_domain}; free-endolysin (top-ranked of "
                         f"{len(cands)} candidate(s))")
        # Generalised JD007 safeguard: a weak free pick (NlpC/P60-only or no
        # CHAP/Amidase and not explicitly endolysin) is cross-checked by tBLASTn.
        weak = ("nlpc" in pick.inferred_domain.lower()
                or not any(t in pick.inferred_domain.lower() for t in ("chap", "amidase", "lysk")))
        if weak and run_tblastn_flag and reference is not None:
            ident, note, suffix = run_tblastn(_record_cache[acc], reference, identity_threshold)
            pick.tblastn_identity, pick.tblastn_note = ident, note
            if suffix == "tblastn-recovered":
                msgs.append(f"[verify] {accession}: keyword pick is weak "
                            f"({pick.inferred_domain}); {note}")
        return pick, msgs

    # No free endolysin by keyword → tBLASTn fallback (requirement 4)
    vaph = [c for c in cands if c.classification == "VAPH"]
    if run_tblastn_flag and reference is not None and acc in _record_cache:
        ident, note, suffix = run_tblastn(_record_cache[acc], reference, identity_threshold)
        msgs.append(f"[{suffix}] {accession}: {note}")
        if vaph:
            v = max(vaph, key=lambda c: c.length)
            v.selected = True
            v.tblastn_identity, v.tblastn_note = ident, note
            v.evidence = f"VAPH ({v.inferred_domain}); no free endolysin; {note}"
            return v, msgs
        return None, msgs

    if vaph:
        v = max(vaph, key=lambda c: c.length)
        v.selected = True
        v.evidence = f"VAPH ({v.inferred_domain}); no free endolysin found by keyword"
        msgs.append(f"[VAPH] {accession}: only a virion-associated hydrolase "
                    f"annotated; run tBLASTn (--run-tblastn) to seek a free endolysin.")
        return v, msgs

    msgs.append(f"[no-endolysin] {accession}: no lysis CDS classified as a free "
                f"endolysin; run tBLASTn (--run-tblastn) with a LysK reference.")
    return None, msgs


# Module-level cache so the selector can reach the SeqRecord for tBLASTn.
_record_cache: dict[str, SeqRecord] = {}


def run_tblastn_helper(acc: str, reference: Path, threshold: float):
    """Thin wrapper that looks up the cached record before running tBLASTn."""
    if acc not in _record_cache:
        return None, "record unavailable", "tblastn-error"
    return run_tblastn(_record_cache[acc], reference, threshold)


# ===========================================================================
# Batch driver
# ===========================================================================

def run(input_dir: Path, fasta_out: Path, csv_out: Optional[Path],
        reference_override: Optional[Path], run_tblastn_flag: bool,
        identity_threshold: float) -> tuple[list[Candidate], list[Candidate]]:
    """Process all genomes. Returns (selected_per_genome, all_candidates)."""
    gb_files = sorted(f for f in input_dir.iterdir()
                      if f.is_file() and f.suffix.lower() in VALID_EXTENSIONS)
    if not gb_files:
        log.error(f"No GenBank files found in '{input_dir}'.")
        sys.exit(1)
    log.info(f"Found {len(gb_files)} GenBank file(s) in '{input_dir}'")

    reference = None
    if run_tblastn_flag:
        reference = build_reference(input_dir, reference_override)
        if find_tblastn() is None:
            log.warning("  --run-tblastn set but BLAST+ (tblastn) is not on PATH; "
                        "commands will be printed instead of executed.")

    all_candidates: list[Candidate] = []
    selected: list[Candidate] = []
    no_candidate: list[str] = []

    for gb in gb_files:
        try:
            record = SeqIO.read(str(gb), "genbank")
        except Exception as exc:
            log.warning(f"  Skipped '{gb.name}': {exc}")
            continue
        _record_cache[record.id.split(".")[0]] = record

        cands = collect_candidates(record)
        all_candidates.extend(cands)
        pick, msgs = select_for_genome(cands, record.id, reference,
                                       run_tblastn_flag, identity_threshold)
        for m in msgs:
            log.info("  " + m)
        if pick is not None:
            selected.append(pick)
            tag = "" if pick.classification == "free-endolysin" else f"  [{pick.classification}]"
            log.info(f"  {gb.name:<18} → {pick.product[:34]} ({pick.length} aa){tag}")
        else:
            no_candidate.append(record.id)
            log.info(f"  {gb.name:<18} → no representative endolysin in annotation")

    # --- Write combined FASTA (selected, annotation-derived sequences) ---
    fasta_out.parent.mkdir(parents=True, exist_ok=True)
    ordered = sorted(selected, key=lambda c: c.accession)
    with open(fasta_out, "w", encoding="utf-8") as fh:
        for c in ordered:
            fh.write(f">{c.fasta_header}\n{c.sequence}\n")
    log.info(f"\nCombined FASTA written: '{fasta_out}'  ({len(ordered)} sequences)")
    if len(ordered) <= INTERPRO_MAX_SEQUENCES:
        log.info(f"  ✓ {len(ordered)} ≤ {INTERPRO_MAX_SEQUENCES}: paste the whole file into InterPro at once.")
    else:
        log.warning(f"  ⚠ {len(ordered)} > {INTERPRO_MAX_SEQUENCES}: split into ≤100-sequence chunks.")

    # --- Write audit CSV (requirement 6) ---
    if csv_out is not None:
        csv_out.parent.mkdir(parents=True, exist_ok=True)
        cols = ["Phage", "Accession", "Protein_ID", "Product", "Length_aa",
                "Inferred_Domain", "InterPro_Domain", "Classification",
                "Selected", "tBLASTn_Identity_pct", "tBLASTn_Note",
                "Runtime_Flag", "Evidence"]
        with open(csv_out, "w", newline="", encoding="utf-8") as fh:
            w = csv.writer(fh)
            w.writerow(cols)
            for c in sorted(all_candidates, key=lambda x: (x.accession, not x.selected)):
                w.writerow([
                    c.organism, c.accession, c.protein_id, c.product, c.length,
                    c.inferred_domain, "",   # InterPro_Domain filled after InterPro
                    c.classification, "Yes" if c.selected else "No",
                    "" if c.tblastn_identity is None else f"{c.tblastn_identity:.1f}",
                    c.tblastn_note, c.runtime_flag, c.evidence,
                ])
        log.info(f"Audit CSV written:      '{csv_out}'  ({len(all_candidates)} candidate rows)")

    if no_candidate:
        log.info("  Genomes with no annotation-derived endolysin (resolve via tBLASTn):")
        for acc in no_candidate:
            log.info(f"    • {acc}")

    return selected, all_candidates


# ===========================================================================
# CLI
# ===========================================================================

def build_parser() -> argparse.ArgumentParser:
    p = argparse.ArgumentParser(
        prog="S4_endolysin_extractor_for_interpro.py",
        description=("Classify and extract endolysins (free vs VAPH vs non-lytic) "
                     "from Staphylococcus phage GenBank records, with a tBLASTn "
                     "fallback, into a combined FASTA for InterPro plus an audit CSV."),
        formatter_class=argparse.RawDescriptionHelpFormatter,
        epilog="""
WORKFLOW (Table 2)
------------------
  1. python S4_endolysin_extractor_for_interpro.py -i GenBank \\
         -o results/endolysin_candidates.faa --csv results/endolysin_audit.csv
  2. Open endolysin_candidates.faa, COPY ALL, paste into InterPro
     (https://www.ebi.ac.uk/interpro/search/sequence/; Pfam, CDD, SUPERFAMILY, Gene3D).
     Fill the InterPro_Domain column of the CSV from the result.
  3. Inspect the CSV: each candidate's Classification and Evidence show why it
     was kept or rejected; 'Selected=Yes' marks the one chosen per genome.

CLASSIFICATIONS
  free-endolysin     : standalone lytic endolysin (validate in InterPro)
  VAPH               : virion-associated (tail/baseplate) hydrolase, not free
  non-lytic          : NADAR/YbiA etc. — rejected as endolysin
  intron-split       : lysK.1 moiety (MN047438, MF398190) — 209 aa N-terminal fragment
  hnh-disrupted      : Sb1M_6168 / Sb1M_9832 — 141 aa CHAP fragment (tBLASTn)
  tblastn-recovered  : free LysK missed by keyword, found by tBLASTn (Maine, JD007)
  divergent-endolysin: divergent free endolysin (e.g., EW NAGPA glycosidase)

tBLASTn FALLBACK
  Add --run-tblastn to enable. Requires NCBI BLAST+ (tblastn) on PATH and a LysK
  reference (auto-extracted from MN336261/Sb1_8383 if present, or --reference).
  Without BLAST+, the exact command is printed and validated identities (from
  the manuscript) are reported.
        """,
    )
    p.add_argument("--input_dir", "-i", type=Path, required=True, metavar="DIR",
                   help="Directory of GenBank files (.gb/.gbk/.gbff)")
    p.add_argument("--output", "-o", type=Path,
                   default=Path("endolysin_candidates.faa"), metavar="FILE",
                   help="Combined FASTA output. Default: endolysin_candidates.faa")
    p.add_argument("--csv", type=Path, default=None, metavar="FILE",
                   help="Audit CSV output (one row per candidate).")
    p.add_argument("--reference", type=Path, default=None, metavar="FILE",
                   help="LysK reference FASTA for tBLASTn (else auto from MN336261).")
    p.add_argument("--run-tblastn", action="store_true",
                   help="Enable the tBLASTn fallback (needs BLAST+ on PATH).")
    p.add_argument("--identity-threshold", type=float, default=90.0, metavar="PCT",
                   help="tBLASTn %% identity above which a hit is a recovered free "
                        "endolysin (else divergent/uncertain). Default: 90.")
    return p


if __name__ == "__main__":
    args = build_parser().parse_args()
    selected, allc = run(args.input_dir, args.output, args.csv,
                         args.reference, args.run_tblastn, args.identity_threshold)

    sep = "=" * 100
    print(f"\n{sep}")
    print("TABLE 2 — SELECTED ENDOLYSIN PER PHAGE (representative; validate in InterPro)")
    print(sep)
    print(f"{'Accession':<13} {'Len':>6}  {'Classification':<20} {'tBLASTn':>8}  {'Product'}")
    print("-" * 100)
    for c in sorted(selected, key=lambda x: x.accession):
        ident = "" if c.tblastn_identity is None else f"{c.tblastn_identity:.0f}%"
        print(f"{c.accession:<13} {c.length:>6}  {c.classification:<20} {ident:>8}  {c.product[:40]}")
    print(sep)
    n_free = sum(1 for c in selected if c.classification == "free-endolysin")
    print(f"  Selected: {len(selected)}  (free-endolysin: {n_free}, "
          f"special/validated: {len(selected) - n_free})")
    print(f"  FASTA : {args.output}")
    if args.csv:
        print(f"  CSV   : {args.csv}   (fill InterPro_Domain after the InterPro run)")
    print(f"\n  Classifications to watch: VAPH (not a free endolysin), non-lytic (rejected),")
    print(f"  intron-split / hnh-disrupted / tblastn-recovered (see CSV Evidence + tBLASTn columns).")
    if not args.run_tblastn:
        print(f"\n  Tip: add --run-tblastn (with BLAST+) to actively recover keyword-missed")
        print(f"       free endolysins (e.g., Maine, JD007) against the Sb1_8383 LysK reference.")
```
