## Supplementary material for "In silico characterization of lysis and host-recognition modules in *Staphylococcus aureus* bacteriophage genomes": https://github.com/HivanA98/In-silico-characterization-of-lysis-and-host-recognition-modules-in-S.-aureus-bacteriophage-genomes: Supplementary_Code_Documentation.docx

**Bioinformatic Data-Extraction Pipeline**

Molecular Characterization of Lytic Bacteriophages Against Resistant Staphylococcus aureus Based on NCBI GenBank Sequences: A Bioinformatic Literature Review

### **1. Manuscript Contribution Map**

Each script generates data for a specific table or figure. Consult this map before running any script.

| **Script** | **Table / Figure** | **Columns Generated** |
| --- | --- | --- |
| S1_genome_statistics.py | TABLE 1 (complete) | Phage, Accession, Class, Family, SubFamily, Genome Size, GC%, CDS Count, tRNA Count, NCBI Status |
| S2_holin_tailfiber_annotation.py | TABLE 2 (partial) | Holin Present, Tail Fiber / RBP Present |
| S3_terl_extractor.py | FIGURE 1 (input) | TerL multi-FASTA -> MAFFT -> MEGA 12.1.2 |
| S4_endolysin_extractor_for_interpro.py | TABLE 2 (domain columns via InterPro) | Endolysin Gene Product, Endolysin Length, Catalytic Domains, Wall Binding Domain |

Table 2 is produced by two scripts: S2 (Holin and Tail Fiber/RBP columns) and S4 followed by InterPro (all domain architecture columns).

### **2. Dependencies (Windows only)**

All scripts run on Windows 10/11 and have been tested with the exact versions below. No dependencies beyond these are required. Script outputs are plain-text CSV and FASTA files.

| **Dependency** | **Version** | **Type / Notes** |
| --- | --- | --- |
| Python | 3.12.10 | Interpreter |
| biopython | 1.87 | pip install — GenBank parsing (all scripts) |
| pandas | 3.0.3 | pip install — CSV output (S1, S2) |
| numpy | 2.4.6 | INDIRECT — auto-installed by pandas; not imported directly |
| MAFFT | web server | https://mafft.cbrc.jp/alignment/server/ (strategy L-INS-i); no local install |
| MEGA | 12.1.2 | External tool — phylogenetics (S3 workflow, Step 3) |
| InterPro | 108.0 | https://www.ebi.ac.uk/interpro/search/sequence/ |

**Installation (Command Prompt or PowerShell)**

pip install biopython==1.87 pandas==3.0.3

numpy 2.4.6 is installed automatically as a pandas dependency; it is not imported directly by any script. All other modules used (argparse, logging, pathlib, dataclasses, sys, typing) are part of the Python 3.12.10 standard library and require no installation.

MAFFT (Windows): https://mafft.cbrc.jp/alignment/software/windows.html

MEGA 12.1.2: https://www.megasoftware.net/

### **3. Input File Preparation**

All scripts require a directory of NCBI GenBank flat files (.gb, .gbk, .gbff). Each file must contain exactly one complete genome record. Name each file by its accession number (e.g., NC_047722.gb).

**Download via Entrez Direct (optional helper, Command Prompt)**

efetch -db nuccore -id NC_047722 -format genbank > GenBank\NC_047722.gb

entrezpy/Entrez Direct is optional and only used for downloading; records may instead be downloaded manually from the NCBI website (Send to > File > GenBank).

### **4. Script Reference**

#### **S1 — Genome Statistics -> TABLE 1 (complete)**

Extracts all Table 1 columns in a single run, including Class, Family, and SubFamily. These three ranks are read from record.annotations['taxonomy'] by ICTV rank suffix: Class from '-viricetes' (e.g., Caudoviricetes), Family from '-viridae' (e.g., Herelleviridae), SubFamily from '-virinae' (e.g., Twortvirinae). This corrects an earlier version that mislabelled the '-viridae' family rank as Class.

python S1_genome_statistics.py -i GenBank -o results\Table1.csv

GUARANTEED NO 'N/A': Some NCBI lineages omit the family rank — the Azeredovirinae phages EW (NC_007056) and SA13 (NC_021863) carry no '-viridae' token. For these, Family is resolved from a documented FAMILY_BY_SUBFAMILY map; subfamilies that NCBI/ICTV place in no family resolve to 'Unassigned' (a valid family-incertae-sedis status, NOT a data error). Verified: Twortvirinae->Herelleviridae; Rakietenvirinae->Rountreeviridae; Azeredovirinae->Unassigned. All Class values are Caudoviricetes for this tailed-phage dataset.

#### **S2 — Holin and Tail Fiber/RBP -> TABLE 2 (partial)**

Detects Holin and Tail Fiber/Receptor-Binding Protein from CDS product annotations. This script intentionally does NOT detect endolysin — that data comes from S4 + InterPro. This focused design reflects the manuscript usage, in which only the Holin Present and Tail Fiber/RBP Present columns of Table 2 derive from text annotation.

python S2_holin_tailfiber_annotation.py -i GenBank -o results\Table2_holin_rbp.csv

Design note on Class/SubFamily placement: Class and SubFamily belong to Table 1, so they are produced by S1 (which already reads the full record), not by S2. This keeps each script's output aligned with a single table and avoids merging Table 1 and Table 2 fields in one file.

**Validation correction (RBP keyword list)**

A comparison against the pre-refactor output revealed that 5 genomes (MN336261, MN336262, MN336263, NC_047725, NC_047726) were incorrectly flagged Tail Fiber/RBP = Yes. The cause was an over-broad keyword 'tail tube protein' added during refactoring. The tail tube is a structural DNA-conduit present in essentially all tailed phages and is NOT a receptor-binding / host-recognition protein; it has been REMOVED from the keyword list. Genuine RBP terms (tail fiber, tail spike, receptor-binding protein, host specificity protein, baseplate receptor-binding, adsorption protein) are retained.

To make every call verifiable, S2 now emits two audit columns — Holin_Evidence and RBP_Evidence — showing the matched keyword and the product annotation text behind each Yes (or a dash for No). These columns let a reviewer confirm each detection against the source GenBank record, and can be dropped when assembling the final Table 2.

#### **S3 — TerL Extraction -> FIGURE 1**

Extracts terminase large subunit (TerL) sequences and writes one combined multi-FASTA file for alignment and phylogenetics.

**TerL Annotation History (documented for reproducibility)**

The initial extraction script failed to identify TerL in 9 of 22 genomes. Investigation revealed two distinct causes, both resolved here:

Case A — Kayvirus group (7 genomes: NC_047722, NC_047723, NC_047724, NC_047725, NC_047726, NC_047727, EU418428): TerL is present (605 aa) but annotated with the 3-letter product name "Ter" instead of "terminase large subunit". Standard keyword search misses this. FIX: an exact-match check (product == "Ter"/"ter") catches all 7 genomes.

Case B — Portland (MT926124) and vB_SauP-436A1 (MN150710): These genomes genuinely lack an annotated TerL. Portland has only "putative encapsidation protein" (415 aa); vB_SauP-436A1 has "DNA packaging protein" (415 aa). Both are micro-phages (~17–18 kb) that do not follow standard Myovirus TerL annotation. DECISION: correctly excluded from phylogenetic analysis and from the output FASTA.

| **Mechanism** | **Check Applied** | **Catches** |
| --- | --- | --- |
| 1. Keyword match | Substring of TERL_KEYWORDS in product qualifier (lowercased) | Standard NCBI TerL annotations (e.g., Twort) |
| 2. Exact product match | product qualifier EXACTLY EQUALS 'Ter' or 'ter' | Kayvirus group — 7 genomes (the V1 fix) |

**Methods Statement (incorporated into the manuscript)**

"Staphylococcus phage Portland (MT926124) and vB_SauP-436A1 (MN150710) were excluded from phylogenetic analysis due to the absence of annotated terminase large subunit sequences, consistent with their atypical small genome sizes (<20 kb) relative to the remaining dataset."

**Complete Figure 1 Workflow**

**Step 1 — Run S3:**

python S3_terl_extractor.py -i GenBank -o results\TerL_combined.faa

**Step 2 — Multiple sequence alignment with MAFFT (web server, no local install):**

```
Open https://mafft.cbrc.jp/alignment/server/
  a. Upload TerL_combined.faa (or paste the FASTA content)
  b. Advanced settings -> strategy: L-INS-i
     (Very slow; recommended for <200 sequences with one
      conserved domain and long gaps; 2 iterative cycles only)
  c. Submit
  d. Save the 'Fasta format' result as TerL_aligned.faa
```

**Step 3 — Phylogenetic tree in MEGA 12.1.2:**

```
Open results\TerL_aligned.faa in MEGA 12.1.2
Phylogeny > Construct/Test Maximum Likelihood Tree
  Substitution model    : LG+G+I
  Rates among sites     : Gamma + Invariant (G+I)
  Bootstrap replicates  : 1000
  Site coverage cutoff  : 80% (Partial Deletion)
  Outgroup              : Staphylococcus phage EW (NC_007056.1)
  Condense tree at      : 50% bootstrap
```

#### **S4 — Endolysin Candidates for InterPro -> TABLE 2 (domain validation)**

Selecting an endolysin by the first product-name keyword match is unsafe. Manual InterPro + tBLASTn validation of this dataset exposed three failures: Maine (MN045228) first-matched a non-lytic N-glycosidase YbiA-like protein, while the real endolysin is a free LysK recovered by tBLASTn at 99% identity; JD007 (NC_019726) first-matched a 295-aa virion-associated NlpC/P60 protein, while the real endolysin is a free 495-aa LysK (tBLASTn 99%, missed by keyword); and Twort (NC_007021) first-matched a 1269-aa phage tail lysozyme, which is a virion-associated peptidoglycan hydrolase (VAPH), not a free endolysin.

The revised script therefore (1) collects ALL lysis-keyword CDS and ranks them; (2) infers a domain class from the annotation and flags non-lytic hits such as NADAR/YbiA — but NOT plain glycosidase, because the NAGPA phosphodiester glycosidase of phage EW is a genuine divergent endolysin; (3) separates free endolysins from virion-associated (VAPH) enzymes; (4) runs an automatic tBLASTn fallback against a LysK reference (auto-extracted from Sb1_8383 / MN336261, or supplied with --reference) when no free endolysin is found by keyword; (5) flags the known intron-split (MN047438, MF398190) and HNH-disrupted (MN336262, MN336263) ORFs; and (6) writes an audit CSV (one row per candidate: matched product, inferred domain, classification, selected flag, tBLASTn identity and coordinates, evidence) alongside the combined FASTA.

Honest scope of domain validation: a GenBank flat file carries product names, not Pfam/InterPro assignments, so true domain assignment still comes from the InterPro step on the FASTA this script produces. Offline, the script infers a domain from the annotation text and length and flags each candidate; the CSV InterPro_Domain column is filled from the InterPro result. tBLASTn is wired as a real tblastn subprocess when NCBI BLAST+ is on PATH; otherwise the exact command is printed and the manuscript-validated identities are reported. The validated Table 2 classifications (InterPro + tBLASTn) are encoded in a KNOWN_CASES table so the script output matches the published table, while any genome not listed is handled by the general heuristic.

Classifications emitted: free-endolysin, VAPH, non-lytic, intron-split, hnh-disrupted, tblastn-recovered, and divergent-endolysin.

python S4_endolysin_extractor_for_interpro.py -i GenBank -o results\endolysin_candidates.faa

MANDATORY FINAL VALIDATION: Open endolysin_candidates.faa, copy ALL text, and paste into InterPro (https://www.ebi.ac.uk/interpro/search/sequence/). Select databases Pfam, CDD, SUPERFAMILY, Gene3D. InterPro identifies CHAP, Amidase, SH3b, LysM, NlpC/P60 domains — the validated content of the Catalytic Domains and Wall Binding Domain columns of Table 2.

Fragmented endolysins (tBLASTn, NOT InterPro): Sb1M_6168 (MN336262) and Sb1M_9832 (MN336263) carry an HNH endonuclease inserted into the endolysin ORF. No intact 490-500 aa endolysin exists; a 57-aa LysM protein and a 166-aa HNH endonuclease are annotated separately. tBLASTn (query = Sb1_8383 endolysin) confirms a 141-aa LysK-homologous region (c29476-29051), consistent with HNH insertion disrupting the ORF (Kornienko et al., 2023). Complete these two Table 2 rows from the tBLASTn result; the script flags both genomes automatically.

**Cross-Validation of Table 1 (taxonomic re-validation)**

The same InterPro submission also re-validates the Table 1 classification. Endolysin domain architecture is family- and subfamily-diagnostic: the CHAP + Amidase_2 + SH3b tri-domain (LysK-type) is characteristic of Twortvirinae / Herelleviridae, so its presence corroborates that assignment; the divergent endolysin content of the Azeredovirinae phages (EW, SA13) is consistent with their separate placement (Family = Unassigned) and outgroup role in Figure 1. Concordance between InterPro domains and the NCBI lineage strengthens BOTH tables; discordance flags a record for manual taxonomic review. A single InterPro run therefore completes Table 2 AND provides an orthogonal check on the Table 1 taxonomy derived independently by S1.

#### **Reconciliation with Primary Literature (vB_SauM-515A1, MN047438)**

Two reports on this phage (Kornienko et al., 2020, Sci Rep; and Viruses, 'Transcriptional Landscape') give values that differ from this toolkit. Both differences are methodological, not code errors: the scripts read the deposited GenBank annotation verbatim, whereas the papers used de-novo / transcriptomic methods invisible to the static DNA annotation.

**(1) CDS/ORF count: 236 (toolkit) vs 238 (papers)**

S1 counts feature.type=='CDS' in the GenBank record as deposited (236). The papers re-annotated with RAST, a de-novo gene-caller, reporting 238 ORFs. Different gene-callers and curation choices give slightly different counts for the same accession; the 2-feature gap is not a counting error and cannot be matched without re-running RAST (a different methodology). MN047438's deposited record also carries no 'complete' keyword, so S1's text-based status inference labels it Draft/Partial. Recommendation: report CDS as the GenBank-deposit count and add a method footnote.

**(2) Endolysin: 209-aa amidase fragment vs full intron-split LysK**

GenBank annotates the endolysin as 'lysK.1', explicitly the 'N-terminal moiety' of LysK (a 209-aa amidase fragment). The full functional LysK (~495 aa: CHAP + Amidase + SH3b) is interrupted by a self-splicing intron, so the N-terminal (lysK.1) and C-terminal (SH3b-bearing) halves are annotated as separate features. The intact enzyme is reconstructed at the RNA level — shown transcriptomically for vB_SauM-515A1 by Kornienko et al. (2020, Viruses, transcription unit TU16) — and is invisible to the static CDS annotation. The same 'lysK.1' architecture appears in vB_SauM-fRuSau02 (MF398190). S4 captures the annotated fragments and auto-flags both accessions; InterPro on a single fragment cannot return the full length. Recommendation for Table 2: pair the InterPro domains (Amidase + SH3b) with a note that the ORF is intron-split and 209 aa is the N-terminal moiety, not the complete protein. This is mechanistically distinct from the HNH-disrupted endolysins (Sb1M_6168, Sb1M_9832), which require tBLASTn (Kornienko et al., 2023).

### **5. Complete Analysis Workflow**

```
REM ---- TABLE 1 (complete, single run) ----
python S1_genome_statistics.py -i GenBank -o results\Table1.csv

REM ---- TABLE 2 (part 1: Holin + Tail Fiber/RBP) ----
python S2_holin_tailfiber_annotation.py -i GenBank -o results\Table2_holin_rbp.csv

REM ---- TABLE 2 (part 2: domain architecture via InterPro) ----
python S4_endolysin_extractor_for_interpro.py -i GenBank -o results\endolysin_candidates.faa
REM  then open the file, copy ALL, paste into https://www.ebi.ac.uk/interpro/

REM ---- FIGURE 1 (TerL -> MAFFT -> MEGA 12.1.2) ----
python S3_terl_extractor.py -i GenBank -o results\TerL_combined.faa
REM  MAFFT web (L-INS-i): https://mafft.cbrc.jp/alignment/server/ -> TerL_aligned.faa
REM  then open TerL_aligned.faa in MEGA 12.1.2 (LG+G+I, 1000 bootstrap)
```

### **6. Changes from Original Scripts**

| **Original Script** | **Issue Identified** | **Resolution** |
| --- | --- | --- |
| Genome Size, GC, CDS, tRNA.py | Single-file only; hardcoded local path | Batch directory processing via argparse (S1) |
| phastest_batch.py | Only Holin + Tail Fiber output used in manuscript | Refocused as S2 (Holin + Tail Fiber only) |
| extract_endolysin.py | Extracted ALL CDS proteins -> InterPro rejects >100 sequences | Replaced by S4: endolysin-candidates only, ONE combined FASTA, paste-ready |
| ekstrak_TerL.py (V1) | Missed 9 genomes: 7 Kayvirus (product='Ter') + Portland + vB_SauP-436A1 | Kayvirus fixed via exact product match; the 2 micro-phages correctly excluded |
| ekstrak_TerL.py (V2, working) | Per-genome FASTA output; required manual concatenation before MAFFT | Merged into S3 with combined multi-FASTA output |
| All scripts | No error handling; hardcoded local Windows paths | try/except + logging; all paths via argparse |
